# The IMD and JNK pathways drive the functional integration of the immune and circulatory systems of mosquitoes

**DOI:** 10.1101/2022.01.26.477938

**Authors:** Yan Yan, Leah T. Sigle, David C. Rinker, Tania Y. Estévez-Lao, John A. Capra, Julián F. Hillyer

**Author notes:** Contributed equally (Yan Yan, Leah T. Sigle).

## Abstract

The immune and circulatory systems of animals are functionally integrated. In mammals, the spleen and lymph nodes filter and destroy microbes circulating in the blood and lymph, respectively. In insects, immune cells that surround the heart valves (ostia), called periostial hemocytes, destroy pathogens in the areas of the body that experience the swiftest hemolymph (blood) flow. An infection recruits additional periostial hemocytes, amplifying heart-associated immune responses. Although the structural mechanics of periostial hemocyte aggregation have been defined, the genetic factors that regulate this process remain less understood. Here, we conducted RNAseq in the African malaria mosquito, *Anopheles gambiae*, and discovered that an infection upregulates multiple components of the IMD and JNK pathways in the heart with periostial hemocytes. This upregulation is greater in the heart with periostial hemocytes than in the circulating hemocytes or the entire abdomen. RNAi-based knockdown then showed that the IMD and JNK pathways drive periostial hemocyte aggregation and alter phagocytosis and melanization on the heart, thereby demonstrating that these pathways regulate the functional integration between the immune and circulatory systems. Understanding how insects fight infection lays the foundation for novel strategies that could protect beneficial insects and harm detrimental ones.

## 1. Introduction

Insect immune cells, called hemocytes, produce pattern recognition receptors that detect microbial invaders, activate immune signaling pathways, and kill pathogens via phagocytosis, lysis, melanization and other mechanisms (1–3). These hemocytes exist in a dynamic body cavity called the hemocoel, where the flow of hemolymph constantly circulates them throughout the body. However, not all hemocytes circulate; one quarter of hemocytes are attached to tissues and remain sessile (4). A sub-population of these sessile hemocytes concentrates on the outer surface of the heart and, more specifically, in the regions surrounding heart valves called ostia (5, 6). These heart-associated hemocytes, called periostial hemocytes, reside in the locations of the body that experience the highest hemolymph flow, and intensely phagocytose bacteria and malaria parasites within seconds of their entry into the hemocoel. As this is happening, additional hemocytes leave circulation and aggregate in the periostial regions of the heart, thereby augmenting the heart-associated immune response (5, 6). Although the kinetics and structural mechanics of periostial immunity have only been described in the African malaria mosquito, *Anopheles gambiae* (5–9), heart-associated immune responses have also been reported in other mosquito species, fruit flies, stick insects, and moths (10–12). Further analysis of insects from 16 different orders showed that the functional integration between the immune and circulatory systems is conserved across the entire insect lineage (13).

In mosquitoes, pattern recognition receptors in the thioester-containing protein family, together with adhesion and phagocytosis factors in the Nimrod protein family, positively regulate immune responses on the heart (14, 15). However, heart-associated immune responses are triggered by both bacterial and malarial infections, and by both peptidoglycan and B-1,3-glucan (5, 6). Hence, the regulatory networks that drive periostial hemocyte aggregation undoubtedly extend beyond pattern recognition receptors and adhesion molecules. In insects, immune responses are primarily driven by pathways such as Toll, immune deficiency (IMD), c-Jun N-terminal kinase (JNK), and Jak/Stat (1, 16). The Toll pathway acts via the transcription factor Rel1, and primarily responds to Gram(+) bacteria, fungi and rodent malaria parasites, whereas the IMD and interlinked JNK pathways act via the transcription factors Rel2 and JNK, respectively, and primarily respond to Gram(–) bacteria and human malaria parasites (17, 18). In fruit flies, the Toll pathway is also involved in hemocyte differentiation, and its constitutive activation disrupts sessile hemocytes (19, 20). The IMD pathway has not been directly linked to cellular immunity but its canonical activator, the peptidoglycan recognition protein PGRP-LC, positively regulates phagocytosis of *Escherichia coli* but not *Staphylococcus aureus* (21). The Jak/Stat pathway acts via the Stat family of transcription factors. In mosquitoes, this pathway responds to malaria parasites and viruses (22, 23), and in fruit flies it controls hemocyte maturation and differentiation (24, 25).

The goal of this study was to identify genes and regulatory pathways that drive periostial hemocyte aggregation in *A. gambiae*. Using an unbiased RNA sequencing (RNAseq) approach that was followed by empirical testing with RNA interference (RNAi), we uncovered that the IMD and JNK pathways drive periostial hemocyte aggregation and immune responses on the heart, thereby regulating the functional integration between the immune and circulatory systems of mosquitoes.

## 2. Results

### 2.1. Infection upregulates immune genes in periostial hemocytes

To identify genes that may drive periostial hemocyte aggregation, we undertook an unbiased RNAseq approach where we sequenced the transcriptome of tissues from mosquitoes that were naïve, injured, or infected with Gram(–) GFP-*E. coli* or Gram(+) *S. aureus* (Figure 1A-B; *Dataset* S1). Three tissues were isolated at 4 hr after treatment: (i) the heart containing periostial hemocytes, (ii) the hemolymph containing circulating hemocytes, and (iii) the entire abdomen. Mosquitoes were assayed at 4 hr after treatment because the number of periostial hemocytes approximately doubles within the first hour of infection, and plateaus by 4 hr after infection (5). In these experiments, most mosquitoes were used for sequencing, but a subset of mosquitoes was used to confirm that infection induces heart-associated cellular immune responses. Indeed, a resident population of periostial hemocytes was present in both naïve and injured mosquitoes, and the number of periostial hemocytes increased 1.8- and 1.6-fold after GFP-*E. coli* and *S. aureus* infection, respectively (Figure 1B-C).

**Figure 1.**
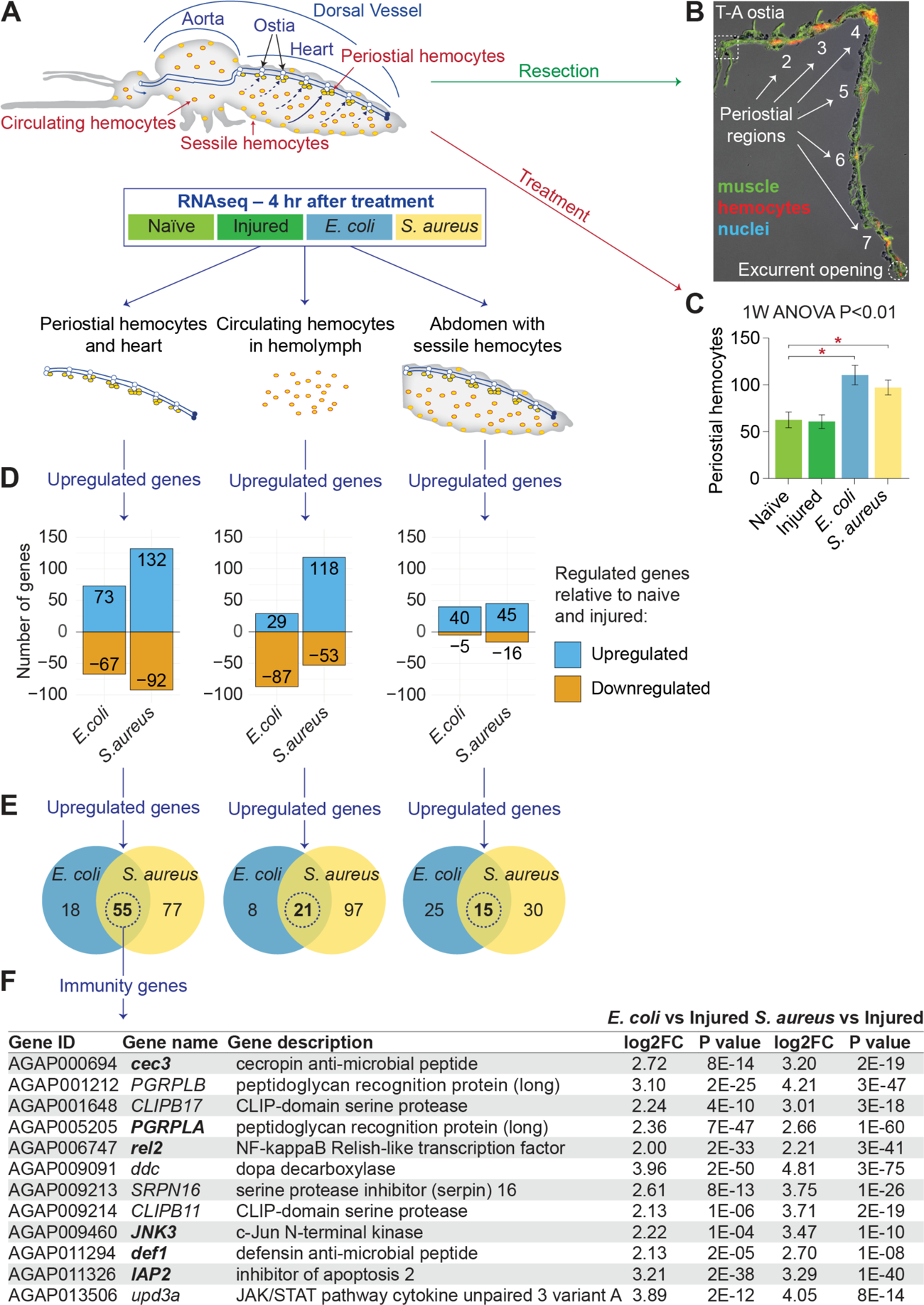
Infection upregulates immune genes in periostial hemocytes. (*A*) The heart with periostial hemocytes, the circulating hemocytes, and the abdomen of mosquitoes that were naïve, injured, or infected with GFP-*E. coli* or *S. aureus* were sequenced by RNAseq at 4 hr post-treatment. (*B*) Fluorescence and DIC overlay shows that the periostial hemocytes (red) remain attached to a resected heart (green). Marked are the periostial regions for each abdominal segment, the thoraco-abdominal (T-A) ostia and the posterior excurrent opening. Image is modified from Sigle and Hillyer (7), and reproduced according to Creative Commons Attribution License CC BY. (*C*) Naïve and injured mosquitoes have resident periostial hemocytes, but infection for 4 hr with GFP-*E. coli* or *S. aureus* induces the aggregation of additional hemocytes at the periostial regions. Whiskers show the standard error of the mean (S.E.M.; n = 14 for all groups). (*D*) Bar plots show the number of genes significantly upregulated or downregulated at 4 hr after GFP-*E. coli* or *S. aureus* infection in the periostial hemocytes and heart (left), the circulating hemocytes (middle), and the entire abdomen (right). (*E*) Venn diagrams show that 55, 21 and 15 genes are significantly upregulated in the periostial hemocytes, the circulating hemocytes or the abdomen, respectively, during both GFP-*E. coli* and *S. aureus* infections. (F) Table listing the immune genes that are among the 55 genes that were upregulated in the heart with periostial hemocytes following both GFP-*E. coli* or *S. aureus* infection. Genes that participate in the IMD and JNK pathway are in bold.

RNAseq revealed that, relative to the heart and periostial hemocytes of naïve and injured mosquitoes, 73 and 132 genes are upregulated more than 4-fold (P < 0.05) in the heart and periostial hemocytes of mosquitoes infected with GFP-*E. coli* and *S. aureus*, respectively (Figure 1D). Because periostial hemocyte aggregation is a dynamic immune response that occurs during any bacterial infection, we hypothesized that its core molecular drivers must be preferentially upregulated in periostial hemocytes following both *E. coli* and *S. aureus* infection. We found that 55 genes met this overlap criterion (Figure 1E; Table S1). Comparing the heart and periostial hemocyte RNAseq data to RNAseq data from (i) circulating hemocytes and (ii) the entire abdomen then revealed that infection upregulates more genes in the heart with periostial hemocytes than in the circulating hemocytes or the abdomen (Figure 1D-E; Tables S1-S3).

We then scrutinized the reported function of the 55 candidate genes. Twelve have immune function (Figure 1F), and strikingly, 6 are part of IMD and JNK pathway: the peptidoglycan recognition protein *PGRP-LA* (26), the inhibitor of apoptosis family member *IAP2* (27), the IMD pathway transcription factor *rel2* (28, 29), the antimicrobial peptides *cec3* (29) and *def1* (30, 31), and the JNK pathway transcription factor *JNK3* (32, 33). Multiple aspects of the IMD and JNK pathway are represented: upstream components such as *PGRP-LA* and *IAP2*, and downstream elements from both branches of the pathway, with the classical IMD cascade represented by *rel2*, *cec3* and *def1*, and the bifurcation into the JNK pathway represented by *JNK3* (27, 34, 35) (Figure S1). Relevant to this finding, activating the IMD pathway induces the expression of the scavenger and phagocytosis receptor *eater* (36), which positively regulates hemocyte adhesion in fruit flies and periostial hemocyte aggregation in mosquitoes (15, 37). The IMD pathway also induces the expression of the thioester containing protein *TEP1*, which is involved in phagocytosis and periostial hemocyte aggregation (14). Moreover, transglutaminases are pleiotropic enzymes that in *Drosophila* inhibit the IMD pathway (38, 39), and a mosquito transglutaminase – *TGase3* – negatively regulates periostial hemocyte aggregation and has infection-dependent effects on the heart rate (40, 41). Finally, the JNK pathway is associated with cell adhesion and phagocytosis by regulating the cytoskeleton (32, 42). The JNK pathway also induces the expression of the engulfment receptor *draper* (43) and *TEP1* (33), both of which affect periostial hemocyte aggregation in mosquitoes (14, 15). All six IMD and JNK pathway genes identified in our unbiased screen were more highly upregulated in the heart and periostial hemocytes than in the other tissues (Figure S2). The remaining six candidate genes with immune function that were identified in our screen – *CLIPB17*, *DDC*, *SRPN16*, *CLIPB11*, *PGRP-LB* and *upd3k* – do not participate in a common pathway.

Because the downregulation of genes could also impact hemocyte aggregation, we assessed the genes that were downregulated after both *E. coli* and *S. aureus* infection, relative to naïve and injured mosquitoes. No genes met this criterion for the abdomen, but 29 genes met this criterion for the heart with periostial hemocytes and 19 genes for the circulating hemocytes (Tables S4 and S5). Five of these genes were shared between the heart with periostial hemocytes and the circulating hemocytes, but none of the 43 genes have a classical role in immunity or carry a function suggestive of involvement in hemocyte aggregation. Therefore, based on the RNAseq analyses we hypothesized that the main driver of periostial hemocyte aggregation is the IMD and JNK pathway.

### 2.2. The IMD pathway positively regulates periostial hemocyte aggregation

To determine whether the IMD pathway regulates periostial hemocyte aggregation, we synthesized double-stranded RNA (dsRNA) to target the IMD cascade transcription factor, *rel2*, and its negative regulator, *caspar*, and achieved RNAi-based silencing of 43% and 37%, respectively, compared to the ds*bla(Ap^R^)* control mosquitoes (Figure S3). Moreover, because *Tep1* is (i) activated by the Rel2 arm of the IMD pathway (44, 45) and (ii) a positive driver of periostial hemocyte aggregation (14), we measured its mRNA abundance in *rel2* and *caspar* RNAi mosquitoes. Silencing of *rel2* did not alter *TEP1* mRNA abundance but silencing of *caspar* increased it. Because *cec3* and *def1* are also activated by Rel2 (29–31), we measured their mRNA abundance in RNAi mosquitoes. Silencing of *rel2* decreased *cec3* and *def1* mRNA abundance whereas silencing of *caspar* increased them (Figure S3). Overall, these data are consistent with the role that the Rel2 arm of the IMD pathway plays in regulating *TEP1*, *cec3* and *def1* expression.

To assess whether the IMD pathway drives heart-associated immune responses, we compared the number and activity of periostial hemocytes in *rel2* and *caspar* RNAi mosquitoes, relative to ds*bla(Ap^R^)* control mosquitoes (Figure 2 and 3). As expected, infection for 4 hr induced the aggregation of hemocytes at the periostial regions of ds*bla(Ap^R^)* mosquitoes, and this aggregation remained in place at 24 hr. In uninfected mosquitoes, knocking down *rel2* did not change the number of periostial hemocytes, but at 4 hr following infection, knockdown of *rel2* decreased the number of periostial hemocytes by 18%, relative to ds*bla(Ap^R^)* mosquitoes (Figure 2A-B). At 24 hr post-infection, the effect of *rel2* RNAi on periostial hemocyte aggregation was diminished. When we instead knocked down *caspar*, the number of periostial hemocytes increased by 39% in uninfected mosquitoes, and infection for 4 and 24 hr increased the number of periostial hemocytes by 30% and 19%, respectively, relative to similarly treated ds*bla(Ap^R^)* mosquitoes. This shows that *rel2* positively regulates periostial hemocyte aggregation whereas *caspar* negatively regulates this process.

**Figure 2.**
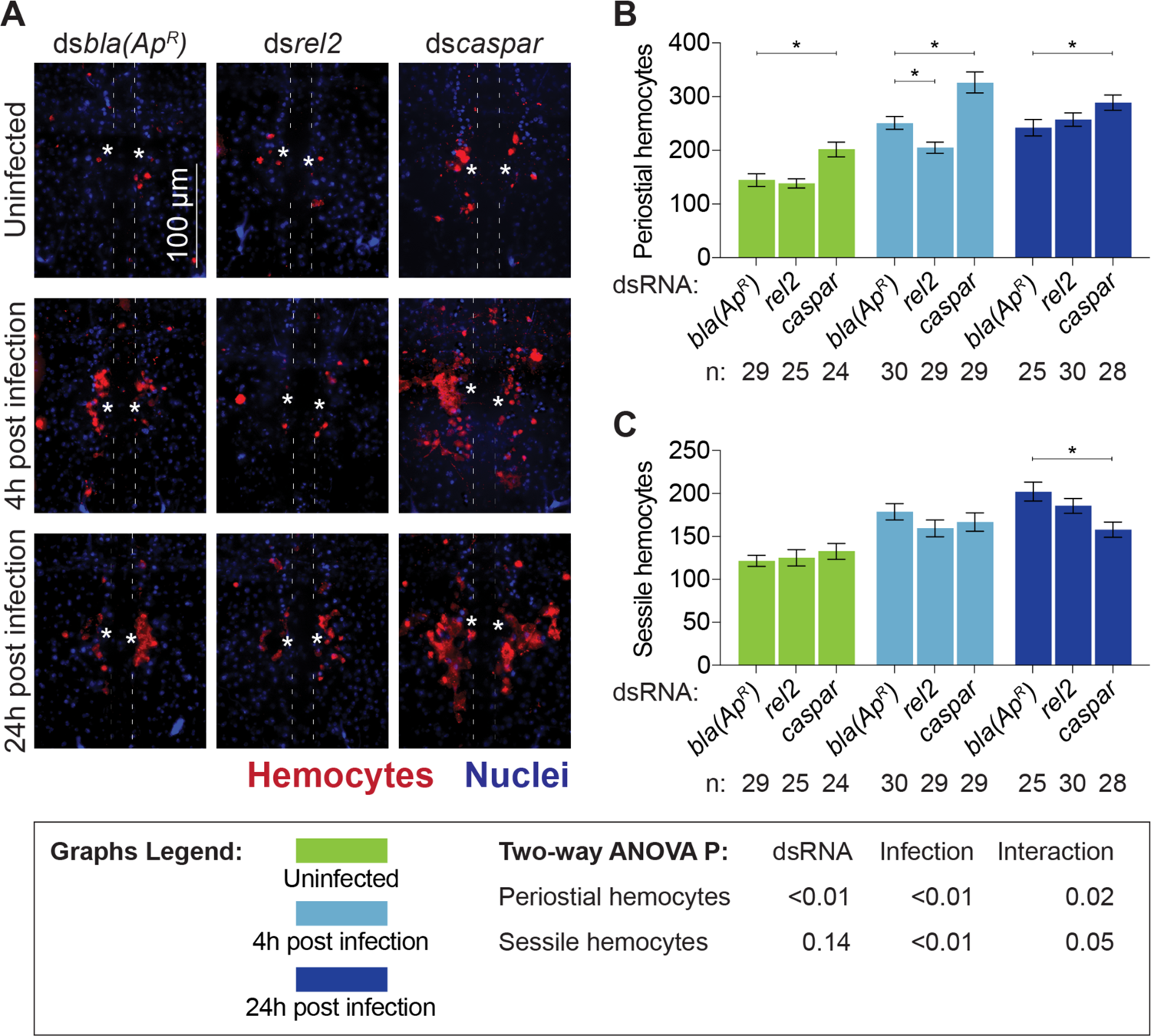
The IMD pathway drives periostial hemocyte aggregation. (*A*) Fluorescence images show periostial hemocytes (CM-DiI; red) surrounding a single pair of ostia (asterisks) on a segment of the heart (outlined by dotted lines) of ds*bla(Ap^R^)*, ds*rel2* and ds*caspar* mosquitoes that were not infected or had been infected with GFP-*E. coli* for 4 or 24 hr. Anterior is on top. (*B-C*) Graphs for ds*bla(Ap^R^)*, ds*rel2* and ds*caspar* mosquitoes that were not infected or had been infected with GFP-*E. coli* for 4 or 24 hr. The graphs show: (*B*) average number of periostial hemocytes; and (*C*) average number of sessile hemocytes outside of the periostial regions in the tergum of abdominal segments 4 and 5. Graphs show the mean and S.E.M. The data were analyzed by two-way ANOVA (bottom box), followed by Dunnett’s multiple comparison test. N indicates sample size. Asterisks in graphs indicate post-hoc P < 0.05.

**Figure 3.**
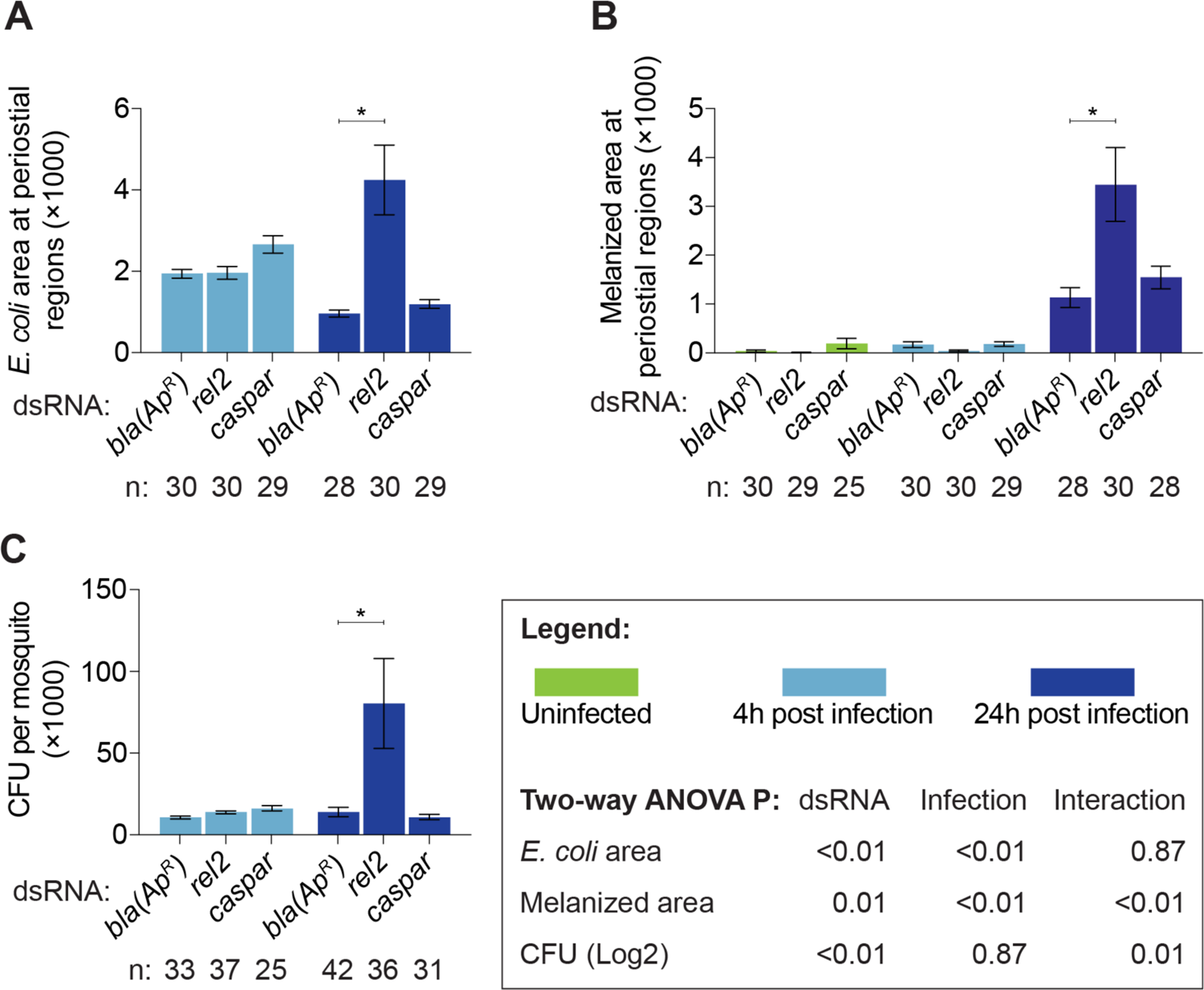
The IMD pathway modulates immune responses on the heart and the systemic antimicrobial response. (*A-C*) Graphs for ds*bla(Ap^R^)*, ds*rel2* and ds*caspar* mosquitoes that were not infected or had been infected with GFP-*E. coli* for 4 or 24 hr. The graphs show: (*A*) pixel area of GFP-*E. coli* in the periostial regions; (*B*) pixel area of melanin in the periostial regions; and (*C*) the systemic GFP-*E. coli* infection intensity. Graphs show the mean and S.E.M. The data were analyzed by two-way ANOVA (bottom box), followed by Dunnett’s multiple comparison test. N indicates sample size. Asterisks in graphs indicate post-hoc P < 0.05.

To determine whether the IMD pathway specifically affects periostial hemocytes and not sessile hemocytes in general, we counted the number of sessile hemocytes outside of the periostial regions in the tergum of abdominal segments 4 and 5 in the same mosquitoes examined for periostial hemocytes (Figure 2C). Knockdown of *rel2* or *caspar* did not alter the number of non-periostial sessile hemocytes in uninfected mosquitoes or mosquitoes infected for 4 hr. However, at 24 hr following infection, *caspar* RNAi decreased non-periostial sessile hemocytes by 22%. These results demonstrate that the IMD pathway regulates periostial hemocyte aggregation while having a minimal effect on the rest of the sessile hemocytes.

Periostial hemocytes phagocytose bacteria, leading to their accumulation at the periostial regions (5, 6). To determine whether the IMD pathway affects the phagocytic activity of periostial hemocytes, we measured the GFP*-E. coli* fluorescence pixel area in the periostial regions of infected mosquitoes (Figure 3A; Figure S4). At 4 hr following infection, the GFP-*E. coli* pixel area was similar regardless of dsRNA treatment. At 24 hr after infection, this area decreased in ds*bla(Ap^R^)* and ds*caspar* mosquitoes, indicating that periostial hemocytes efficiently destroyed the pathogens. However, at 24 hr following infection, *rel2* RNAi significantly increased GFP-*E. coli* accumulation at the periostial regions. Periostial hemocytes also phagocytose melanized bacteria (5, 6), and one melanization related-gene – dopa decarboxylase or *DDC* (46, 47) – was upregulated in both the heart with periostial hemocytes and the circulating hemocytes. Therefore, we measured melanin accumulation in the periostial regions (Figure 3B; Figure S4). Melanin was absent in uninfected mosquitoes and mosquitoes that were infected for 4 hr. However, at 24 hr following infection, *rel2* RNAi increased melanin deposits at the periostial regions whereas *caspar* RNAi had no effect.

The increased accumulation of GFP-*E. coli* and melanin in *rel2* RNAi mosquitoes at 24 hr after infection could be due to either (i) enhanced phagocytosis by periostial hemocytes, or (ii) higher bacterial proliferation in the hemocoel, which places increased pressure on the phagocytosis response. To differentiate between these two scenarios, we measured the systemic GFP-*E. coli* infection intensity and observed that, at 4 hr after infection, the bacterial intensity was similar for all dsRNA treatments, but at 24 hr, knockdown of *rel2* resulted in a higher infection intensity than treatment with ds*caspar* or ds*bla(Ap^R^)* (Figure 3C). This suggests that *rel2* is essential for proper bacterial killing in the hemocoel, and therefore, knocking it down increases infection intensity in a manner that leads to increased phagocytosis in the periostial regions. However, because silencing *rel2* and *caspar* had opposite effects on periostial hemocyte aggregation at 4 hr, a time when dsRNA treatment does not impact infection intensity, we conclude that the IMD pathway is a positive regulator of periostial hemocyte aggregation.

PGRP-LC is the canonical activator of the IMD pathway (1), but PRGP-LA (i) activates the IMD pathway in the midgut of mosquitoes and in the barrier epithelia of fruit flies (26, 48), (ii) is expressed in *Drosophila* hemocytes (49), and (iii) is upregulated in the heart and periostial hemocytes (Figure 1F). Therefore, we tested the involvement of PGRP-LA in periostial hemocyte aggregation. *PGRP-LA* has three splice forms, so to knock it down, we synthesized ds*PGRP-LA-RARB* to target the *RA* and *RB* splice forms, and ds*PGRP-LA-RC* to target the *RC* splice form. Using these dsRNAs, we achieved RNAi-based silencing that ranged from 56% to 77%, compared to the ds*bla(Ap^R^)* control mosquitoes (Figure S5). *PGRP-LA* knockdown with any dsRNA did not alter the number of periostial hemocytes, the number of non-periostial sessile hemocytes, melanin accumulation at the periostial regions, or systemic infection intensity (Figure S6). However, knockdown of *PGRP-LA-RC* increased phagocytosis of GFP-*E. coli* in the periostial regions, which suggests that the RC splice form is involved in the phagocytosis response.

### 2.3. The JNK pathway positively regulates periostial hemocyte aggregation

We next tested whether the JNK pathway regulates periostial hemocyte aggregation. *A. gambiae* encodes two JNK genes: *JNK1* and *JNK3* (50). Therefore, we synthesized double-stranded RNA (dsRNA) to target the JNK cascade transcription factors, *JNK1* and *JNK3*, and their negative regulator, *puckered* (*puc*). Because of high sequence identity between *JNK1* and *JNK3*, ds*JNK1/3* simultaneously targeted both JNK genes. RNAi-based knockdown resulted in 33%, 44% and 31% reduction in mRNA abundance of *JNK1*, *JNK3* and *puc*, respectively (Figure S7). Moreover, because the JNK pathway has also been implicated in the expression of *TEP1* (51), we measured the mRNA abundance of *TEP1*, *cec3* and *def1* in *JNK1*/*JNK3* and *puc* RNAi mosquitoes. Manipulating the JNK pathway did not have a clear impact on effector gene expression (Figure S7). The lack of clarity on how the JNK pathway controls the expression of *TEP1*, *cec3* or *def1* could be due to their co-regulation by other pathways, or a consequence of incomplete gene silencing of *JNK1/3* and *puc*.

To assess whether the JNK pathway drives heart-associated immune responses, we compared the number and activity of periostial hemocytes in *JNK1/3* and *puc* RNAi mosquitoes, relative to ds*bla(Ap^R^)* control mosquitoes (Figure 4 and 5). Regardless of infection status, the number of periostial hemocytes was statistically similar between ds*JNK1/3* and ds*bla(Ap^R^)* mosquitoes, although ds*JNK1/3* mosquitoes averaged fewer periostial hemocytes than control mosquitoes for all treatments (Figure 4A-B). Knockdown of *puc*, however, increased the number of periostial hemocytes 1.6-fold in uninfected mosquitoes, and 1.4- and 1.5-fold at 4 and 24 hr following GFP-*E. coli* infection, respectively, relative to similarly treated ds*bla(Ap^R^)* mosquitoes. To further determine whether *puc* specifically affects periostial hemocytes or sessile hemocytes in general, we quantified the number of non-periostial sessile hemocytes on the tergum of the same mosquitoes (Figure 4C). *JNK1/3* RNAi mosquitoes showed a trend of fewer non-periostial sessile hemocytes than control mosquitoes regardless of infection status. Treatment with ds*puc*, however, increased the number of non-periostial sessile hemocytes for all treatments. This demonstrates that the JNK pathway positively regulates both periostial and non-periostial sessile hemocyte abundance.

**Figure 4.**
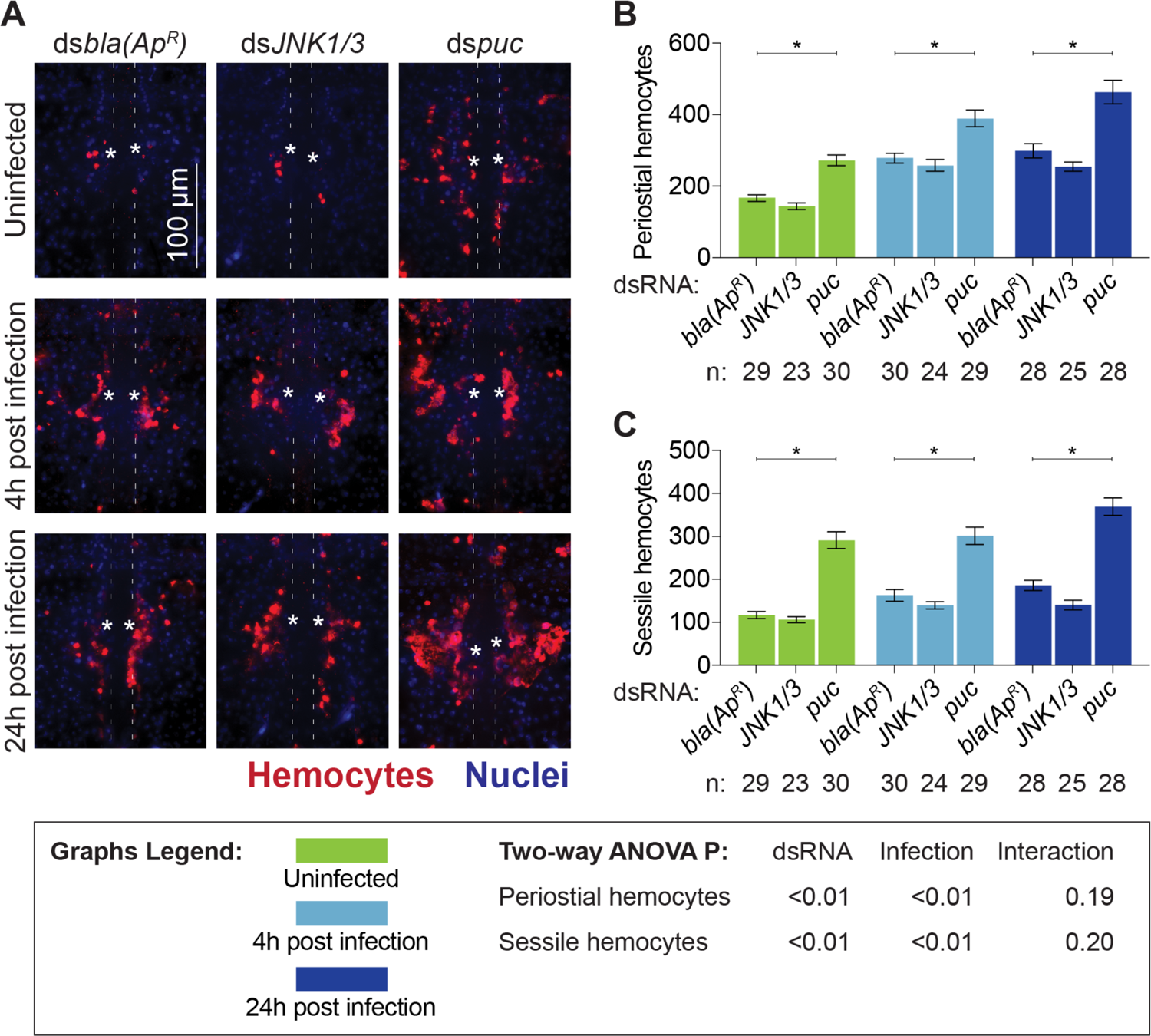
The JNK pathway drives periostial hemocyte aggregation. (*A*) Fluorescence images show periostial hemocytes (CM-DiI; red) surrounding a single pair of ostia (asterisks) on a segment of the heart (outlined by dotted lines) of ds*bla(Ap^R^)*, ds*JNK1/3* and ds*puc* mosquitoes that were not infected or had been infected with GFP-*E. coli* for 4 or 24 hr. Anterior is on top. (*B-C*) Graphs for ds*bla(Ap^R^)*, ds*JNK1/3* and ds*puc* mosquitoes that were not infected or had been infected with GFP-*E. coli* for 4 or 24 hr. The graphs show: (*B*) average number of periostial hemocytes; and (*C*) average number of sessile hemocytes outside of the periostial regions in the tergum of abdominal segments 4 and 5. Graphs show the mean and S.E.M. The data were analyzed by two-way ANOVA (bottom box), followed by Dunnett’s multiple comparison test. N indicates sample size. Asterisks in graphs indicate post-hoc P < 0.05.

**Figure 5.**
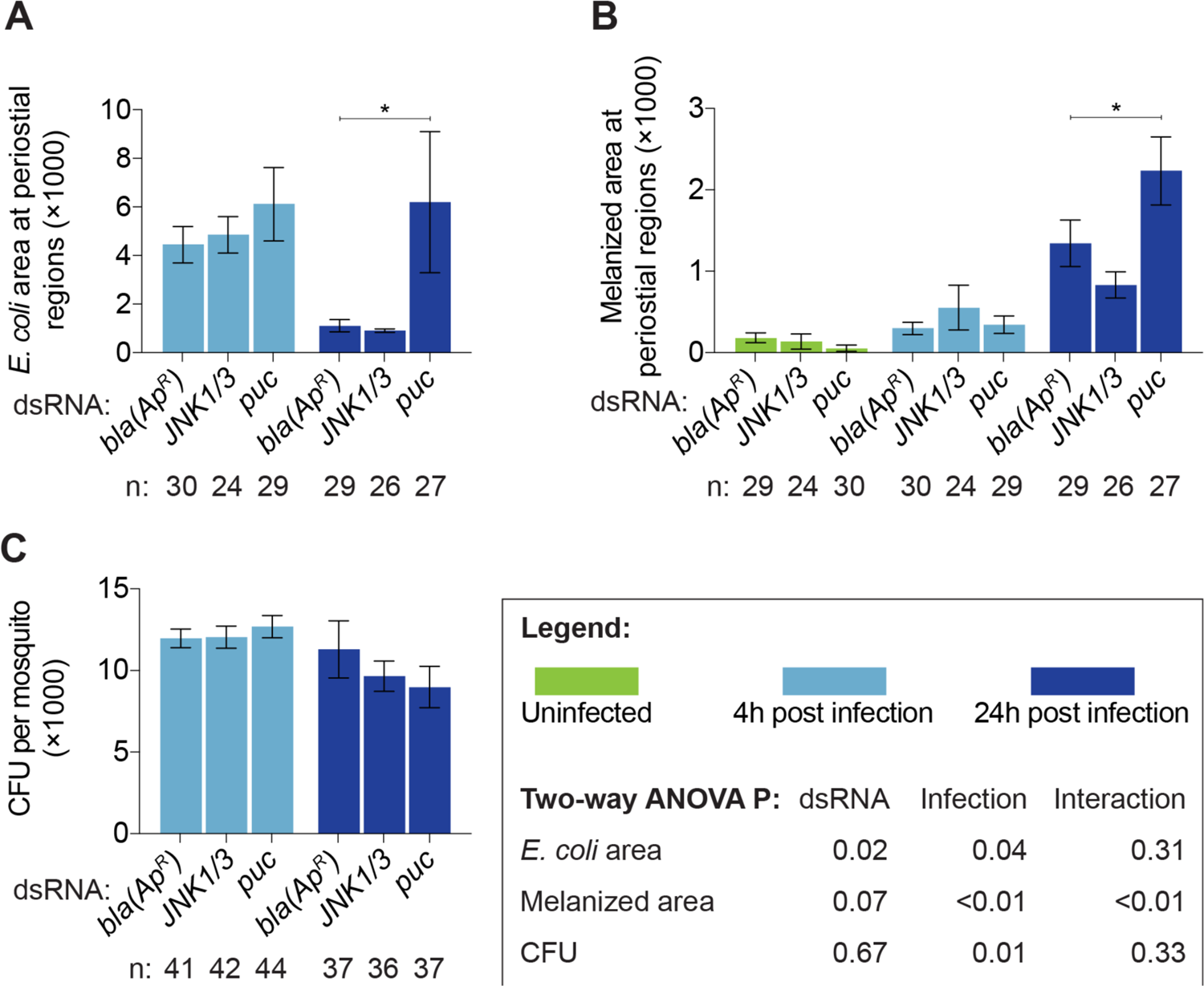
The JNK pathway modulates immune responses on the heart but not the systemic antimicrobial response. (*A-C*) Graphs for ds*bla(Ap^R^)*, ds*JNK1/3* and ds*puc* mosquitoes that were not infected or had been infected with GFP-*E. coli* for 4 or 24 hr. The graphs show: (*A*) pixel area of GFP-*E. coli* in the periostial regions; (*B*) pixel area of melanin in the periostial regions; and (*C*) the systemic GFP-*E. coli* infection intensity. Graphs show the mean and S.E.M. The data were analyzed by two-way ANOVA (bottom box), followed by Dunnett’s multiple comparison test. N indicates sample size. Asterisks in graphs indicate post-hoc P < 0.05.

We next measured whether the JNK pathway affects the phagocytic activity of periostial hemocytes (Figure 5A; Figure S8). At 4 hr following infection, the GFP-*E. coli* pixel area was similar regardless of dsRNA treatment. However, at 24 hr following infection, periostial hemocytes in ds*puc* mosquitoes phagocytosed more GFP-*E. coli* than the ds*bla(Ap^R^)* mosquitoes. We then quantified the melanized bacteria that were sequestered by periostial hemocytes (Figure 5B; Figure S8). Melanin deposits were undetectable in uninfected mosquitoes and mosquitoes at 4 hr post infection; however, at 24 hr following GFP-*E. coli* infection, more melanin was present in ds*puc* mosquitoes than in ds*bla(Ap^R^)* controls.

The difference in GFP-*E. coli* and melanin accumulation in ds*puc* mosquitoes could be due to (i) enhanced phagocytosis by periostial hemocytes, or (ii) higher bacterial proliferation in the hemocoel. To differentiate between these two scenarios, we quantified the systemic *E. coli* infection intensity and found that the bacterial intensity was similar regardless of dsRNA treatment (Figure 5C). Therefore, this suggests that *puc* negatively regulates hemocyte adhesion and phagocytic activity, and demonstrates that the JNK pathway is a positive regulator of periostial immune responses.

## 3. Discussion

Previous microarray and RNAseq analyses revealed genes and signaling pathways that are active in hemocytes (52–58), including those that are activated in specific subpopulations of hemocytes (59–61). However, these studies only focused on circulating hemocytes, and therefore, failed to capture the biology of sessile hemocytes. Yet, one quarter of hemocytes are sessile (4), and they play significant roles in hematopoiesis (62, 63), wound healing (64), and pathogen killing (9, 65, 66). Therefore, extrapolating the molecular signatures of circulating hemocytes to those of sessile hemocytes likely misses the essential factors that make sessile hemocytes conduct their specific immune activities. Periostial hemocytes are a subpopulation of sessile hemocytes that reside on the heart, where they sequester and kill pathogens in areas of high hemolymph flow (5, 6, 13). To better understand how an infection drives the migration of hemocytes to the heart and how these immune cells kill pathogens at the periostial regions, we sequenced the periostial hemocyte transcriptome and discovered that the IMD and JNK pathway drives periostial immune responses (Figure 6).

**Figure 6.**
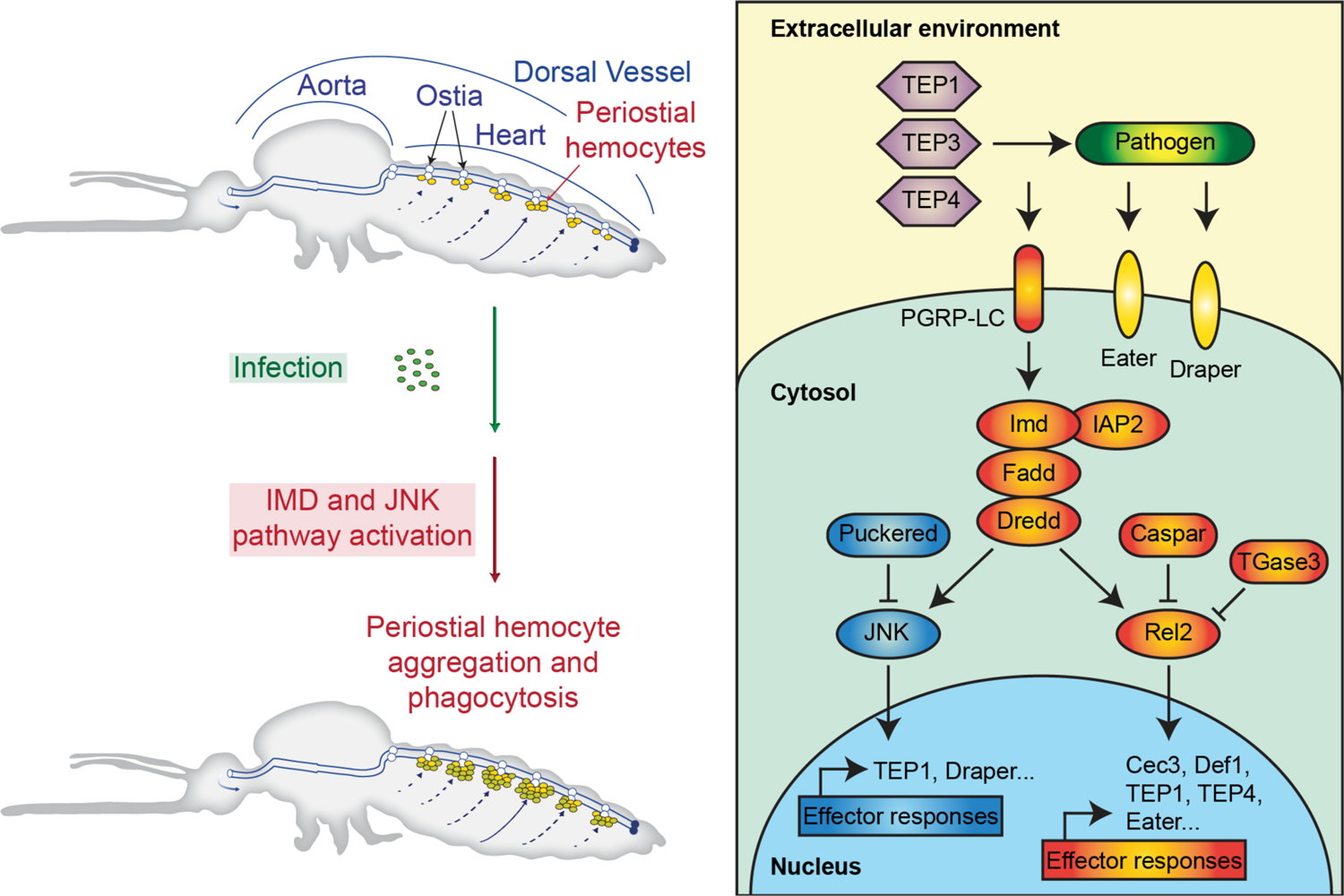
The IMD and JNK pathways, together with TEP proteins (TEP1, TEP3 and TEP4), Nimrod proteins (Draper and Eater) and a transglutaminase, regulate heart-associated immune responses in mosquitoes.

The IMD pathway controls the production of antimicrobial peptides (1, 27), but here we show that the IMD pathway also regulates a cellular immune response: the transition of hemocytes from a circulating to a sessile state on the heart. Specifically, knockdown of the positive regulator of the IMD pathway, *rel2*, decreases infection-induced periostial hemocyte aggregation, whereas knockdown of the negative regulator of the IMD pathway, *caspar*, increases the number of periostial hemocytes. We hypothesize that the IMD pathway drives periostial hemocyte aggregation via two cascades that are also driving phagocytosis: the TEP1-TEP3-LRP1-CED6L and TEP4-BINT2-CED2L-CED5L pathways (67). The IMD pathway induces the expression of *TEP1* and *TEP4* (30, 31, 68), and both of these genes positively regulate periostial hemocyte aggregation (14). Moreover, two downstream molecules – the low-density-lipoprotein-receptor-related protein LRP1 and the beta integrin BINT2 – are transmembrane proteins that in mammals and insects have overlapping functions in phagocytosis and adhesion (69–71), and therefore, likely facilitate the adhesion of hemocytes to the heart. In this study, we also confirmed the positive role that REL2 plays in pathogen killing (29, 44, 68, 72–78). When we systemically knocked down the expression of *rel2*, bacteria proliferated uncontrollably during the later stages of infection. We hypothesize that knocking down *rel2* initially suppresses periostial hemocyte aggregation, but as the infection progresses and the intensity increases, the necessity of periostial immune responses also increases, and this pressure recruits hemocytes to the heart and enhances their collective phagocytic activity. Overall, our data show that the IMD pathway drives periostial hemocyte aggregation during the early stages of infection and limits systemic infection intensity during the later stages of infection.

The JNK pathway modulates mosquito longevity, regulates oviposition, and limits infection with malaria parasites and viruses (33, 51, 79–82). Here, we found that overexpressing the JNK pathway by knocking down the negative regulator, *puc*, increases the number of periostial and adjacent sessile hemocytes. This suggests that the JNK pathway positively regulates hemocyte adhesion. Indeed, the *Drosophila* ortholog of mosquito *JNK1* and *JNK3*, called *basket*, is involved in the formation of actin-rich and focal adhesion kinase-rich placodes in hemocytes (83). The JNK pathway also induces hemocyte differentiation in *Drosophila*, producing large lamellocytes that adhere and encapsulate parasitoid eggs (84). We found that overactivating the JNK pathway does not alter infection intensity in mosquitoes, but it increases the accumulation of *E. coli* and melanin on the heart, suggesting that the JNK pathway positively regulates the phagocytosis and melanization responses. Indeed, JNK1 positively regulates melanization in mosquitoes that are refractory to malaria (51), and overexpressing the JNK pathway in aphids increases the melanin-producing activity of phenoloxidase and the phagocytic activity of hemocytes (85). We hypothesize that the JNK pathway regulates phagocytosis by periostial hemocytes in a manner that involves two proteins already known to be involved in periostial hemocyte aggregation: TEP1 and draper (14, 15). The JNK pathway activates the expression of both of these genes (33, 80, 86), and TEP1 opsonizes pathogens for phagocytosis whereas draper activates phagocytic processes (14, 15, 67, 87–89). In our study, we could not distinguish between the roles that *JNK1* and *JNK3* play in periostial hemocyte aggregation. However, simultaneous knockdown of both resulted in phenotypes that were opposite of *puc* knockdown, strongly suggesting that JNK1 or JNK3 – or both – positively regulates periostial hemocyte aggregation.

Because both the IMD and JNK pathways induce the production of *TEP1* (30, 33, 68, 80), we hypothesize that they share the TEP1-TEP3-LRP1-CED6L phagocytosis cascade that leads to hemocyte aggregation. However, we also hypothesize that the IMD and JNK pathways use additional, independent mechanisms to regulate periostial hemocyte aggregation for three reasons. First, even though the two pathways share upstream signaling molecules, they bifurcate and activate their own set of effector genes (34, 35). Second, knocking down the components of the IMD and JNK pathways resulted in different phenotypes, especially for periostial and non-periostial sessile hemocytes. Third, in *Drosophila*, draper and the BINT2 ortholog (Integrin *βν*) function independently (90), and in mosquitoes, draper is regulated by the JNK pathway whereas BINT2 is involved in the IMD-regulated and TEP4-mediated phagocytosis cascade (31, 67, 70, 86).

Bacteria, malaria parasites, and fungal components all induce periostial hemocyte aggregation, and this process is structurally conserved across the entire insect lineage (5, 6, 13). Therefore, it is likely that other proteins contribute to this process. Indeed, infection also induces the upregulation of *Upd3* by periostial hemocytes, which is a ligand for the receptor of the JAK/STAT pathway (91). The TOLL pathway also induces hemocyte proliferation (92, 93) and the expression of *TEP1* (44), which positively regulates periostial hemocyte aggregation (14). Beyond immune pathways, periostial hemocyte aggregation may also be under the influence of neuronal and hormonal control. For example, injecting mosquitoes with the neuropeptide, allatotropin, increases the number of periostial and non-periostial sessile hemocytes (94). Therefore, the regulation of periostial hemocyte aggregation is not expected to be limited to the IMD and JNK pathways.

To date, we have tested the involvement of several hemocyte-produced factors on periostial hemocyte aggregation. However, the heart itself may produce complementary factors that drive this immune response. For example, a fibroblast growth factor is highly regulated in the heart and periostial hemocytes. Its ortholog in *Drosophila*, called *Branchless*, is expressed in the heart and pericardial cells, and regulates hemocyte differentiation in the lymph glands of the larval heart (95, 96). Moreover, reducing the expression of the *Drosophila* cardiac extracellular matrix (ECM) proteins, *Pericardin* and *Laminin A*, disrupts the formation of the cardiac ECM and lowers the number of hemocytes on the heart (12, 97). These data suggest that the heart and its associated structures facilitate the binding of circulating hemocytes to the periostial regions.

The mosquito genome encodes three transglutaminase genes, and our RNAseq experiment revealed that two of them are upregulated in the heart and periostial hemocytes following infection. In *Drosophila*, transglutaminase negatively regulates the IMD pathway by (i) crosslinking the transcription factor, Relish, into a polymer and (ii) incorporating natural primary amines into the DNA binding site of Relish (38, 39). We recently tested whether transglutaminases are involved in periostial responses in *A. gambiae*, and found that *TGase3* – but not *TGase1* or *TGase2* – negatively regulates periostial hemocyte aggregation during the early stages of infection and the sequestration of melanin by periostial hemocytes during the later stages of infection (40). Moreover, disrupting *TGase2* and *TGase3* has infection-dependent effects on the heart rate (41). Combined, these data further support our conclusion that the IMD pathway is a positive driver of heart-associated immune responses (Figure 6).

Heart-associated immune responses occur in insects from at least 16 different taxonomic orders, including insects that are of agricultural, urban, and medical importance (13). Understanding the genetic factors that drive periostial hemocyte aggregation in *A. gambiae* sheds light on how this medically important insect and other insects of societal importance fight the pathogens that invade them. Indeed, the TEP gene family, the Nimrod gene family (Eater, Nimrod and Draper) and the JNK pathway are conserved amongst insects (50, 98–100), and although some insects in the order Hemiptera lack components of the IMD pathway, all insects queried to date still have a functional IMD-based immune response (101–104). Therefore, we conclude that the IMD and JNK pathways, together with TEP and Nimrod proteins, are primary regulators of the heart-associated immune responses in insects.

## 4. Materials and Methods

### 4.1. Mosquitoes, bacteria, infection, and replication

*Anopheles gambiae,* Giles sensu stricto (G3; Diptera: Culicidae), were maintained at 27 °C, 75% relative humidity, and a 12 hr:12 hr light:dark photoperiod (105). Experiments were done using female adults fed 10% sucrose. After any treatment, mosquitoes of a given treatment were housed together, returned to the environmental chamber, and given access to 10% sucrose. Tetracycline resistant, GFP-expressing *E. coli* (modified DH5*α*; GFP-*E. coli*) and *S. aureus* (RN6390) were grown in Luria-Bertani (LB) and tryptic soy broth, respectively, at 37 °C in a shaking incubator. Dilutions of the bacterial cultures were injected at the thoracic anepisternal cleft using a Nanoject III (Drummond Scientific Company, Broomall, PA). The infection dose was determined by plating the cultures and counting the colony forming units (CFUs).

In this study, a biological replicate is an independent experiment that utilizes mosquitoes from an independent egg batch. A technical replicate is an experimental resampling of a biological replicate. All data collected were included in the analysis; no data were excluded.

### 4.2. RNAseq: treatment, tissue collection, RNA isolation and library preparation

Six-day-old mosquitoes were randomly divided into four groups: (i) naïve (unmanipulated), (ii) injured by injecting 69 nL of sterile LB, (iii) infected by injecting 69 nL of GFP-*E. coli* (∼75,779 CFUs), and (iv) infected by injecting 69 nL of *S. aureus* (∼39,451 CFUs). At 4 hr after treatment, three tissues were isolated: (i) the heart with periostial hemocytes, (ii) hemolymph with circulating hemocytes, and (iii) the entire abdomen. To isolate the heart, mosquitoes were bisected along the coronal plane in RNase-free PBS, and the heart with the periostial hemocytes was resected by severing the alary muscles and detaching it from the cuticle (7). To isolate circulating hemocytes, hemolymph was perfused by making an incision in the last abdominal segment, injecting RNase-free PBS into the hemocoel through the cervical membrane, and collecting the first 2 drops that exited the abdomen (106). The abdomens were isolated by bisecting mosquitoes on the transverse plane along the thoraco-abdominal junction.

To isolate RNA from hearts or abdomens, samples were homogenized in TRIzol Reagent (Invitrogen, Carlsbad, CA), extracted following the TRIzol protocol, and resuspended in Buffer RLT (Qiagen, Hilden, Germany) with 2% 2-mercaptoethanol. The RNA was further purified using the RNeasy Micro Kit (Qiagen), DNase treated, and eluted in RNase-free water. To isolate RNA from circulating hemocytes, perfused hemolymph was collected in Buffer RLT and 2% 2-mercaptoethanol, and RNA was isolated using the RNeasy Micro Kit as above.

No a priori statistical method was used to pre-determine sample sizes. The number of RNAseq samples and biological replicates was determined prior to the initiation of the study, and was based on our determination that three biological replicates is a conservative approach to detect statistical differences (if any) in gene expression. Three biological replicates were conducted; each heart sample contained 72 hearts, each hemolymph sample contained perfusate from 108 mosquitoes, and each abdomen sample contained 36 abdomens.

The integrity and quantity of RNA was assessed on a 2100 Bioanalyzer (Agilent Technologies, Santa Clara, CA) using an RNA 6000 Series II Nano kit for abdomen samples and a Pico kit for the heart and hemolymph samples. The library for sequencing was prepared using 1 µg of RNA and the NEBNext Ultra kit (New England BioLabs, Ipswich, MA), according to the protocol for low-input samples. Library quality and concentration were assessed using a DNA 1000 Series II kit on the 2100 Bioanalyzer.

### 4.3. RNAseq: Illumina sequencing and differential gene expression analysis

All 36 samples – 4 treatments, 3 tissue types and 3 biological replicates – were sequenced across 3 lanes on an Illumina HiSeq 3000 (paired-end, 75 base pair read) at Vanderbilt University’s Vantage facility. Reads were mapped to the *A. gambiae* genome (AgamP4.7) by STAR (107, 108). The number of uniquely mapped reads per sample averaged 23,698,077 (range: 18,487,153 – 31,487,002), which represented 94.03% of the total reads in a sample. Differential expression was calculated based on reads per kilobase per million by DESeq2 (109). Genes were considered significantly regulated at log_2_ fold change ≥ 2 or log_2_ fold change ≤ −2 and the Benjamini–Hochberg adjusted P < 0.05. RNAseq results are presented in Dataset S1, which includes read counts and log2 fold changes. Using four-fold expression difference as the criterion, together with experimental replication and statistical validation, is a conservative approach that is proven to yield accurate and reliable results (110). RNAseq data were deposited into NCBI (https://www.ncbi.nlm.nih.gov) as SRA data PRJNA730047.

### 4.4. Double-stranded RNA (dsRNA) synthesis and RNA interference (RNAi)

Double-stranded RNA was synthesized for *rel2*, *caspar*, *PGRP-LA*, *JNK1* and *JNK3*, and *puc*. *A. gambiae* cDNA was amplified by PCR using gene-specific primers with T7 promoter tags (Table S6). dsRNA was synthesis using the MEGAscript T7 Kit (Applied Biosystems) as described (14, 15). As a negative control, dsRNA was synthesized for the non-mosquito gene, *bla(Ap^R^)*, using DNA from *E. coli* BL21(DE3) containing the pET-46 plasmid as template (EMD Chemicals, Gibbstown, NJ) (14, 15).

Two- or three-day-old mosquitoes were intrathoracically injected 300 ng of dsRNA to initiate systemic gene silencing. Four days later, mosquitoes were divided into two groups for phenotypic analyses: (i) uninfected and (ii) infected with GFP-*E. coli* (∼16,528 CFUs). Injured mosquitoes were not included in the experiments because injury does not induce periostial hemocyte aggregation (4–6). RNAi efficiency of the targeted genes and mRNA abundance of the downstream effector genes *TEP1*, *cec3* and *def1* was determined by qPCR (14). Briefly, RNA was isolated using TRIzol from 10 whole bodies for each time and treatment, purified, and used for cDNA synthesis using the SuperScript III First-Strand Synthesis System for RT-PCR (Invitrogen) as described (14, 15). qPCR was conducted using gene-specific primers (Table S6) and Power SYBR Green PCR Master Mix (Applied Biosystems, Foster City, CA) on a Bio Rad CFX Connect Real-Time Detection System (Hercules, CA). Relative quantification was conducted using the 2^-ΔΔC^_T_ method, with *RpS7* as the reference and *RpS17* as a control (111). Two to three biological replicates were conducted per gene, and the value for each biological replicate is the average of two or three technical replicates.

### 4.5. Fluorescence labeling and mosquito dissection

Hemocytes were labeled with the Vybrant CM-DiI Cell-Labeling Solution (Invitrogen) as we described (5). Briefly, live mosquitoes were injected ∼0.4 µL of 67 µM CM-DiI and 1.08 mM Hoechst 33342 (nuclear stain; Invitrogen) in PBS, incubated at 27 °C for 20 min, and injected 16% paraformaldehyde. Ten min later, abdomens were bisected along a coronal plane, and the dorsal portions containing the heart and periostial hemocytes were mounted on glass slides using Aqua-Poly/Mount (Polysciences; Warrington, PA). CM-DiI stains live hemocytes, and we have used this technique to monitor hemocyte location, number and migration, as well as hemocyte-mediated phagocytosis and melanization (4-9, 13-15, 40).

### 4.6. Microscopy and image acquisition

Specimens were imaged on a Nikon Eclipse Ni-E compound microscope connected to a Nikon Digital Sight DS-Qi1 camera and Advanced Research NIS Elements software (Nikon, Tokyo, Japan). Z-stacks for bright field, red fluorescence (hemocytes), green fluorescence (GFP-*E. coli*) and blue fluorescence (nuclei) were acquired using a linear encoded Z-motor. Specific channels were selected and all images within a stack were combined into a two-dimensional image using the Extended Depth of Focus (EDF) function.

### 4.7. Quantification of hemocytes

Hemocytes, labeled with both CM-DiI and Hoechst 33342, were counted manually by examining all images within a Z-stack (14). A cell was a periostial hemocyte if adjacent to an ostium, and a non-periostial sessile hemocyte if attached to the abdominal wall outside of a periostial region (4, 5). Periostial hemocytes were counted within abdominal segments 2-7 (all abdominal periostial regions) whereas non-periostial sessile hemocytes were only counted on the tergum of segments 4 and 5. Hemocytes were not counted on the aorta, the thoraco-abdominal ostia or the excurrent openings because few hemocytes are there, and infection does not induce aggregation at those locations (7, 8). Data were analyzed by two-way ANOVA, followed by Dunnett’s multiple comparison test, with ds*bla(Ap^R^)* mosquitoes as the reference (Prism 9, GraphPad Software, San Diego, CA). Data for RNAi phenotypic experiments are from individual mosquitoes, which were sampled across three to four biological trials. The exception is experiments for PGRP-LA, where mosquitoes were sampled across two biological trials.

### 4.8. Quantification of GFP-E. coli and melanin at the periostial regions

In NIS-Elements, each periostial region in an EDF image was delineated using the ROI tool. GFP-*E. coli* at the periostial regions was calculated by measuring the area of pixels with intensities above a threshold that distinguished GFP-*E. coli* from background fluorescence (14). Melanin was quantified by measuring the area of pixels with intensities below a threshold that distinguished dark melanized areas from non-melanized areas (14). For each mosquito, measurements from all ROIs were added. Data were analyzed by two-way ANOVA, followed by Dunnett’s multiple comparison test, with ds*bla(Ap^R^)* mosquitoes as the reference.

### 4.9. Quantification of bacterial infection intensity

Mosquitoes that were infected with tetracycline resistant GFP-*E. coli* for 4 or 24 hr were homogenized individually in PBS. A dilution of the homogenate was spread on LB agar containing tetracycline, and plates were incubated overnight at 37 °C. The CFUs were counted and used to calculate infection intensity. Data were analyzed by two-way ANOVA, followed by Dunnett’s multiple comparison test, with ds*bla(Ap^R^)* mosquitoes as the reference. For *rel2* and *caspar* RNAi mosquitoes, data were first log_2_ transformed to achieve normality, and then analyzed by two-way ANOVA.

## Data accessibility

The datasets generated and analyzed during the current study are available in an accompanying supplementary dataset file.

## Author’s contributions

Conceptualized and designed the research: YY, LTS, JFH; Performed the research: YY, LTS, TYE; Contributed resources: JAC, JFH; Analyzed data: YY, LTS, DCR, JFH; Wrote the original draft of the paper: YY, JFH; Administered the project and acquired funding: JFH.

## Acknowledgments

We thank S. Williams, J. Sanner, C. Meier, L. Martin and L. Jabbur for useful discussions and commenting on this manuscript.

## Supplementary tables and figures

**Table S1.** Upregulated genes in the heart with periostial hemocytes at 4 hr following both GFP-*E. coli* and *S. aureus* infection.

| Gene ID | Gene name | Gene description (if annotated) | *E. coli* vs Injured |  | *S. aureus* vs Injured |  |
| --- | --- | --- | --- | --- | --- | --- |
|  |  |  | Log2Foldchange | Adjusted P | Log2Foldchange | Adjusted P |
| AGAP000694 | *cec3* | cecropin anti-microbial peptide | 2.72 | 8E-14 | 3.20 | 2E-19 |
| AGAP000786 |  |  | 2.88 | 4E-06 | 3.12 | 2E-07 |
| AGAP001160 |  |  | 3.03 | 3E-21 | 4.46 | 1E-47 |
| AGAP001212 | *PGRPLB* | peptidoglycan recognition protein (long) | 3.10 | 2E-25 | 4.21 | 3E-47 |
| AGAP001508 |  |  | 3.19 | 4E-07 | 5.03 | 2E-17 |
| AGAP001563 |  | glycerol kinase | 2.58 | 7E-12 | 2.70 | 3E-13 |
| AGAP001597 |  |  | 2.14 | 9E-04 | 2.14 | 6E-04 |
| AGAP001610 |  |  | 4.07 | 2E-21 | 4.13 | 2E-22 |
| AGAP001634 |  |  | 3.52 | 1E-14 | 2.48 | 1E-07 |
| AGAP001648 | *CLIPB17* | CLIP-domain serine protease | 2.24 | 4E-10 | 3.01 | 3E-18 |
| AGAP001828 |  |  | 2.13 | 2E-16 | 2.38 | 9E-21 |
| AGAP002064 |  |  | 2.05 | 7E-18 | 2.29 | 1E-22 |
| AGAP002065 |  | alpha-tocopherol transfer protein | 2.14 | 1E-13 | 2.64 | 3E-21 |
| AGAP002291 |  | ebony | 2.81 | 1E-30 | 3.62 | 2E-51 |
| AGAP002323 |  |  | 2.82 | 2E-11 | 3.31 | 4E-16 |
| AGAP002331 | *esp* | sodium-independent sulfate anion transporter | 3.15 | 8E-20 | 2.94 | 1E-17 |
| AGAP002359 |  | carbonic anhydrase | 2.62 | 2E-02 | 4.62 | 4E-06 |
| AGAP002536 |  |  | 2.48 | 3E-13 | 2.75 | 1E-16 |
| AGAP002883 |  |  | 2.28 | 3E-24 | 2.69 | 5E-34 |
| AGAP003488 |  | nucleotide exchange factor SIL1 | 2.02 | 2E-06 | 2.49 | 1E-09 |
| AGAP004170 |  |  | 2.18 | 4E-21 | 2.73 | 8E-34 |
| AGAP004181 |  | Fibroblast growth factor | 2.53 | 1E-06 | 2.40 | 4E-06 |
| AGAP004826 |  |  | 3.04 | 4E-09 | 4.30 | 2E-18 |
| AGAP005205 | *PGRPLA* | peptidoglycan recognition protein (long) | 2.36 | 7E-47 | 2.66 | 1E-60 |
| AGAP005610 |  |  | 2.44 | 1E-06 | 4.99 | 2E-27 |
| AGAP005839 |  | MFS transporter, FLVCR family | 3.65 | 4E-41 | 3.97 | 3E-49 |
| AGAP006426 |  | cyanogenic beta-glucosidase | 2.65 | 2E-03 | 5.58 | 2E-13 |
| AGAP006670 |  | gamma-glutamyl hydrolase | 2.31 | 8E-24 | 2.90 | 4E-38 |
| AGAP006671 |  |  | 3.18 | 6E-18 | 3.85 | 2E-27 |
| AGAP006674 |  | chymotrypsin-like protease (Precursor) | 2.93 | 2E-02 | 7.46 | 1E-12 |
| AGAP006745 |  |  | 2.35 | 4E-29 | 2.78 | 3E-41 |
| AGAP006747 | *rel2* | NF-kappaB Relish-like transcription factor | 2.00 | 2E-33 | 2.21 | 3E-41 |
| AGAP007659 |  |  | 2.61 | 4E-15 | 3.57 | 1E-28 |
| AGAP008279 | *d7l2* | D7 long form salivary protein | 16.43 | 2E-05 | 13.73 | 3E-04 |
| AGAP008851 |  | E3 ubiquitin-protein ligase mind-bomb | 2.66 | 2E-26 | 2.95 | 8E-33 |
| AGAP009091 | *ddc* | dopa decarboxylase | 3.96 | 2E-50 | 4.81 | 3E-75 |
| AGAP009099 | *TGase3* | Transglutaminase | 3.98 | 2E-17 | 4.05 | 1E-18 |
| AGAP009100 | *TGase1* | Transglutaminase | 3.20 | 3E-37 | 3.76 | 1E-51 |
| AGAP009145 |  |  | 2.12 | 1E-13 | 3.03 | 4E-28 |
| AGAP009213 | *SRPN16* | serine protease inhibitor (serpin) 16 | 2.61 | 8E-13 | 3.75 | 1E-26 |
| AGAP009214 | *CLIPB11* | CLIP-domain serine protease | 2.13 | 1E-06 | 3.71 | 2E-19 |
| AGAP009460 | *JNK3* | c-Jun N-terminal kinase | 2.22 | 1E-04 | 3.47 | 1E-10 |
| AGAP009985 |  | 4-nitrophenylphosphatase | 2.19 | 3E-08 | 3.16 | 4E-17 |
| AGAP010197 |  | SPRY domain-containing SOCS box protein 1 | 2.31 | 2E-20 | 2.85 | 2E-31 |
| AGAP010398 |  | Flavin-containing monooxygenase FMO GS-OX-like 1 | 2.78 | 8E-08 | 5.85 | 1E-33 |
| AGAP010934 |  |  | 2.94 | 1E-30 | 3.17 | 3E-36 |
| AGAP011294 | *def1* | defensin anti-microbial peptide | 2.13 | 2E-05 | 2.70 | 1E-08 |
| AGAP011326 | *IAP2* | inhibitor of apoptosis 2 | 3.21 | 2E-38 | 3.29 | 1E-40 |
| AGAP011588 |  |  | 2.60 | 2E-29 | 3.33 | 4E-49 |
| AGAP011654 |  | membrane dipeptidase | 2.13 | 5E-05 | 3.58 | 3E-13 |
| AGAP011764 |  |  | 2.71 | 5E-63 | 2.87 | 4E-71 |
| AGAP011992 |  | lysosomal acid lipase | 2.72 | 7E-05 | 3.01 | 4E-06 |
| AGAP012166 |  |  | 3.70 | 2E-13 | 2.92 | 8E-09 |
| AGAP013506 | *upd3a* | JAK/STAT pathway cytokine unpaired 3 variant A | 3.89 | 2E-12 | 4.05 | 8E-14 |
| AGAP028615 | *nat5* | nutrient amino acid transporter 5 | 2.54 | 4E-10 | 4.28 | 2E-28 |

**Table S2.** Upregulated genes in the circulating hemocytes at 4 hr following both GFP-*E. coli* and *S. aureus* infection.

| Gene ID | Gene name | Gene description (if annotated) | *E. coli* vs Injured |  | *S. aureus* vs Injured |  |
| --- | --- | --- | --- | --- | --- | --- |
|  |  |  | Log2Foldchange | Adjusted P | Log2Foldchange | Adjusted P |
| AGAP000899 |  |  | 3.31 | 3E-07 | 4.74 | 8E-16 |
| AGAP001212 | *PGRPLB* | peptidoglycan recognition protein (long) | 2.68 | 2E-18 | 4.39 | 4E-51 |
| AGAP001430 |  |  | 2.27 | 2E-18 | 3.39 | 7E-43 |
| AGAP001508 |  |  | 2.06 | 3E-03 | 3.26 | 1E-07 |
| AGAP002291 |  | ebony | 2.46 | 3E-23 | 3.73 | 2E-55 |
| AGAP004115 |  | cystinosin | 2.07 | 5E-03 | 5.38 | 5E-18 |
| AGAP005610 |  |  | 2.45 | 8E-08 | 4.09 | 7E-23 |
| AGAP005639 | *abcb2* | ATP-binding cassette transporter (ABC transporter) family B member 2 | 2.23 | 1E-12 | 3.47 | 1E-31 |
| AGAP007651 |  | growth arrest and DNA-damage-inducible protein | 2.19 | 8E-11 | 3.30 | 2E-25 |
| AGAP009091 | *ddc* | dopa decarboxylase | 2.06 | 2E-13 | 2.93 | 3E-28 |
| AGAP010349 |  |  | 2.28 | 3E-05 | 3.50 | 1E-12 |
| AGAP010387 |  | alanine-glyoxylate aminotransferase | 2.29 | 3E-03 | 3.54 | 3E-07 |
| AGAP010571 |  |  | 2.43 | 5E-06 | 3.25 | 3E-11 |
| AGAP010855 |  | solute carrier family 45, member 1/2/4 | 3.24 | 7E-06 | 5.23 | 7E-16 |
| AGAP011326 | *IAP2* | inhibitor of apoptosis 2 | 2.26 | 2E-18 | 2.58 | 4E-25 |
| AGAP011391 |  |  | 2.07 | 7E-07 | 3.62 | 8E-22 |
| AGAP011588 |  |  | 2.43 | 7E-25 | 3.18 | 8E-45 |
| AGAP011589 |  |  | 2.95 | 2E-02 | 3.46 | 3E-03 |
| AGAP012565 |  |  | 3.58 | 7E-02 | 3.78 | 5E-02 |
| AGAP013285 | *ir7u* | ionotropic receptor IR7u | 2.36 | 3E-03 | 3.46 | 9E-07 |
| AGAP028615 | *nat5* | nutrient amino acid transporter 5 | 2.27 | 9E-08 | 4.06 | 2E-25 |

**Table S3.** Upregulated genes in the abdomen at 4 hr following both GFP-*E. coli* and *S. aureus* infection.

| Gene ID | Gene name | Gene description (if annotated) | *E. coli* vs Injured |  | *S. aureus* vs Injured |  |
| --- | --- | --- | --- | --- | --- | --- |
|  |  |  | Log2Foldchange | Adjusted P | Log2Foldchange | Adjusted P |
| AGAP001212 | *PGRPLB* | peptidoglycan recognition protein (long) | 3.22 | 4E-26 | 3.97 | 8E-41 |
| AGAP001508 |  |  | 2.54 | 3E-04 | 4.63 | 5E-14 |
| AGAP001610 |  |  | 3.33 | 1E-13 | 2.75 | 1E-09 |
| AGAP001648 | *CLIPB17* | CLIP-domain serine protease | 2.04 | 8E-08 | 3.27 | 1E-20 |
| AGAP002291 |  | ebony | 2.52 | 3E-24 | 2.47 | 1E-23 |
| AGAP006670 |  | gamma-glutamyl hydrolase | 2.08 | 1E-18 | 2.33 | 9E-24 |
| AGAP006745 |  |  | 2.06 | 8E-21 | 2.39 | 2E-28 |
| AGAP007651 |  | growth arrest and DNA-damage-inducible protein | 2.17 | 2E-10 | 2.16 | 1E-10 |
| AGAP007659 |  |  | 2.13 | 2E-09 | 2.45 | 7E-13 |
| AGAP009145 |  |  | 2.18 | 2E-13 | 3.51 | 2E-36 |
| AGAP010398 |  | Flavin-containing monooxygenase FMO GS-OX-like 1 | 2.62 | 2E-06 | 5.93 | 3E-33 |
| AGAP010934 |  |  | 2.59 | 6E-23 | 2.25 | 1E-17 |
| AGAP011294 | *def1* | defensin anti-microbial peptide | 2.04 | 2E-04 | 2.39 | 3E-06 |
| AGAP011326 | *IAP2* | inhibitor of apoptosis 2 | 2.16 | 2E-16 | 2.17 | 8E-17 |
| AGAP013348 |  |  | 2.40 | 4E-22 | 3.34 | 2E-43 |

**Table S4.** Downregulated genes in the heart with periostial hemocytes at 4 hr following both GFP-*E. coli* and *S. aureus* infection.

| Gene ID | Gene name | Gene description (if annotated) | *E. coli* vs injury |  | *S. aureus* vs injury |  |
| --- | --- | --- | --- | --- | --- | --- |
|  |  |  | Log2Foldchange | Adjusted P | Log2Foldchange | Adjusted P |
| AGAP000313 |  | alanine-glyoxylate aminotransferase 2-like | -2.19 | 2E-12 | -2.24 | 3E-13 |
| AGAP001009 |  |  | -2.28 | 2E-04 | -3.11 | 1E-05 |
| AGAP001098 |  | DAMOX | -2.25 | 8E-03 | -2.73 | 1E-03 |
| AGAP002429 | *CYP314A1* | cytochrome P450 | -2.46 | 1E-03 | -3.41 | 2E-06 |
| AGAP003494 |  | sugar transporter ERD6-like 7 | -2.75 | 1E-08 | -2.63 | 3E-08 |
| AGAP004248 | *GPXH3* | glutathione peroxidase 3 | -2.32 | 9E-05 | -2.43 | 3E-05 |
| AGAP004954 |  | rhythmically expressed gene 2 protein | -2.10 | 7E-12 | -2.30 | 3E-14 |
| AGAP005498 |  | Phospholipid scramblase | -2.13 | 4E-05 | -2.90 | 4E-09 |
| AGAP005499 |  | dehydrogenase/reductase SDR family member 11 precursor | -2.53 | 4E-04 | -3.33 | 4E-06 |
| AGAP005752 |  | glucosyl/glucuronosyl transferases | -2.29 | 2E-06 | -2.33 | 7E-07 |
| AGAP005753 |  | glucosyl/glucuronosyl transferases | -2.55 | 2E-10 | -2.60 | 9E-11 |
| AGAP006576 |  | malate/L-lactate dehydrogenase | -2.17 | 1E-21 | -2.72 | 1E-33 |
| AGAP008207 | *CYP6Y2* | cytochrome P450 | -2.43 | 2E-33 | -2.63 | 1E-38 |
| AGAP008467 |  | solute carrier family 6 (neurotransmitter transporter, amino acid) member 5/7/ | -3.02 | 2E-17 | -2.74 | 2E-15 |
| AGAP008593 |  |  | -2.47 | 3E-02 | -3.27 | 3E-03 |
| AGAP009194 | *GSTE2* | glutathione S-transferase epsilon class 2 | -2.26 | 4E-15 | -2.02 | 6E-13 |
| AGAP010004 |  |  | -2.13 | 2E-04 | -2.81 | 1E-06 |
| AGAP010367 |  |  | -2.04 | 3E-04 | -2.70 | 2E-06 |
| AGAP010794 |  |  | -2.21 | 3E-07 | -2.71 | 3E-10 |
| AGAP010854 |  | solute carrier family 45, member 1/2/4 | -5.05 | 1E-03 | -3.15 | 3E-02 |
| AGAP012391 |  | DUF1298 domain-containing protein | -2.20 | 2E-04 | -2.63 | 5E-06 |
| AGAP012399 |  | maltase | -2.15 | 6E-07 | -2.51 | 2E-09 |
| AGAP012644 |  |  | -2.72 | 6E-06 | -3.51 | 1E-08 |
| AGAP013094 |  | Elongation of very long chain fatty acids protein | -2.62 | 8E-07 | -2.49 | 2E-06 |
| AGAP013211 |  |  | -2.32 | 6E-03 | -3.79 | 6E-05 |
| AGAP013526 |  |  | -3.31 | 7E-05 | -4.00 | 7E-05 |
| AGAP028049 |  | Carrier domain-containing protein | -2.21 | 1E-09 | -2.13 | 3E-09 |
| AGAP028128 | *ABCC9* | ATP-binding cassette transporter (ABC transporter) family C member 9 | -2.11 | 1E-09 | -2.60 | 4E-13 |
| AGAP028586 |  |  | -3.00 | 6E-09 | -3.46 | 2E-10 |

**Table S5.** Downregulated genes in the circulating hemocytes at 4 hr following both GFP-*E. coli* and *S. aureus* infection.

| Gene ID | Gene name | Gene description (if annotated) | *E. coli* vs injury |  | *S. aureus* vs injury |  |
| --- | --- | --- | --- | --- | --- | --- |
|  |  |  | Log2Foldchange | Adjusted P | Log2Foldchange | Adjusted P |
| AGAP000144 |  | progesterin and adipoQ receptor family member 3 | -2.48 | 1E-03 | -2.80 | 2E-04 |
| AGAP001106 |  | Disconnected protein | -2.32 | 9E-11 | -2.77 | 1E-15 |
| AGAP001476 |  | Innexin inx2 | -2.24 | 1E-09 | -2.81 | 1E-15 |
| AGAP001635 |  | sodium-coupled monocarboxylate transporter 2 | -2.53 | 2E-10 | -2.61 | 3E-11 |
| AGAP002072 |  |  | -2.08 | 4E-05 | -2.59 | 2E-07 |
| AGAP002429 | *CYP314A1* | cytochrome P450 | -2.14 | 7E-03 | -3.53 | 1E-06 |
| AGAP003494 |  | sugar transporter ERD6-like 7 | -2.62 | 4E-07 | -2.68 | 6E-08 |
| AGAP004954 |  | rhythmically expressed gene 2 protein | -2.09 | 3E-11 | -2.67 | 3E-18 |
| AGAP005752 |  | glucosyl/glucuronosyl transferases | -2.06 | 5E-05 | -2.50 | 1E-07 |
| AGAP007990 |  | glucosyl/glucuronosyl transferases | -2.75 | 2E-02 | -3.13 | 6E-03 |
| AGAP008237 |  | Putative G-protein coupled receptor GPCR | -2.01 | 2E-07 | -3.28 | 2E-18 |
| AGAP008629 |  | N-acetyltransferase domain-containing protein | -2.01 | 1E-07 | -3.47 | 8E-16 |
| AGAP009381 |  | Cellular retinaldehyde-binding protein | -2.18 | 9E-05 | -3.37 | 2E-07 |
| AGAP009382 |  |  | -4.31 | 1E-02 | -4.17 | 1E-02 |
| AGAP010794 |  |  | -2.33 | 4E-06 | -3.53 | 4E-10 |
| AGAP011714 |  |  | -2.18 | 4E-08 | -2.28 | 2E-09 |
| AGAP012216 |  | sterol O-acyltransferase | -2.15 | 8E-08 | -3.36 | 1E-16 |
| AGAP028204 |  | Alpha-mannosidase | -2.04 | 1E-02 | -2.04 | 1E-02 |
| AGAP028745 |  |  | -2.03 | 4E-03 | -2.12 | 2E-03 |

**Table S6.**
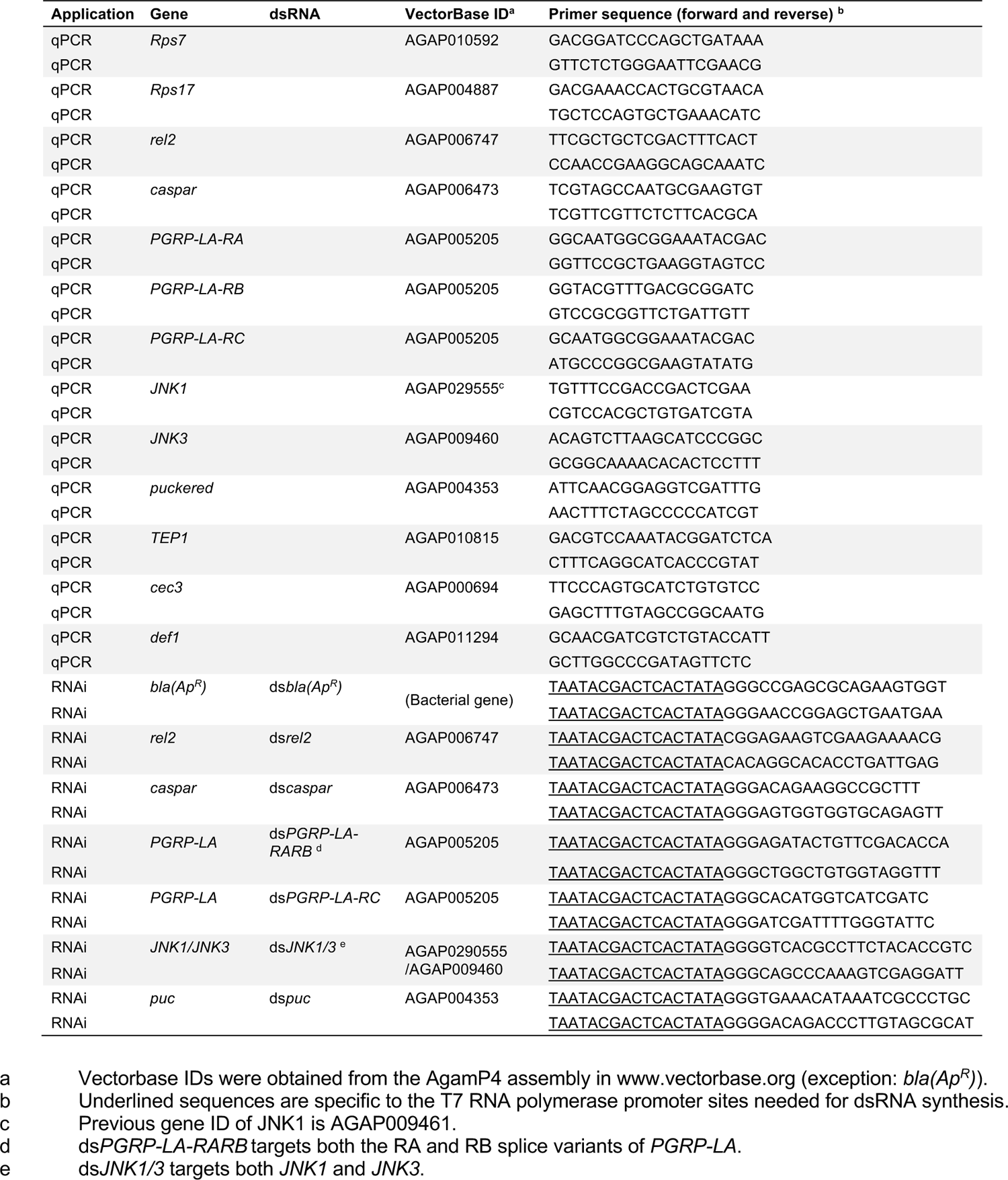
Gene names, gene IDs, and primers used.

**Figure S1.**
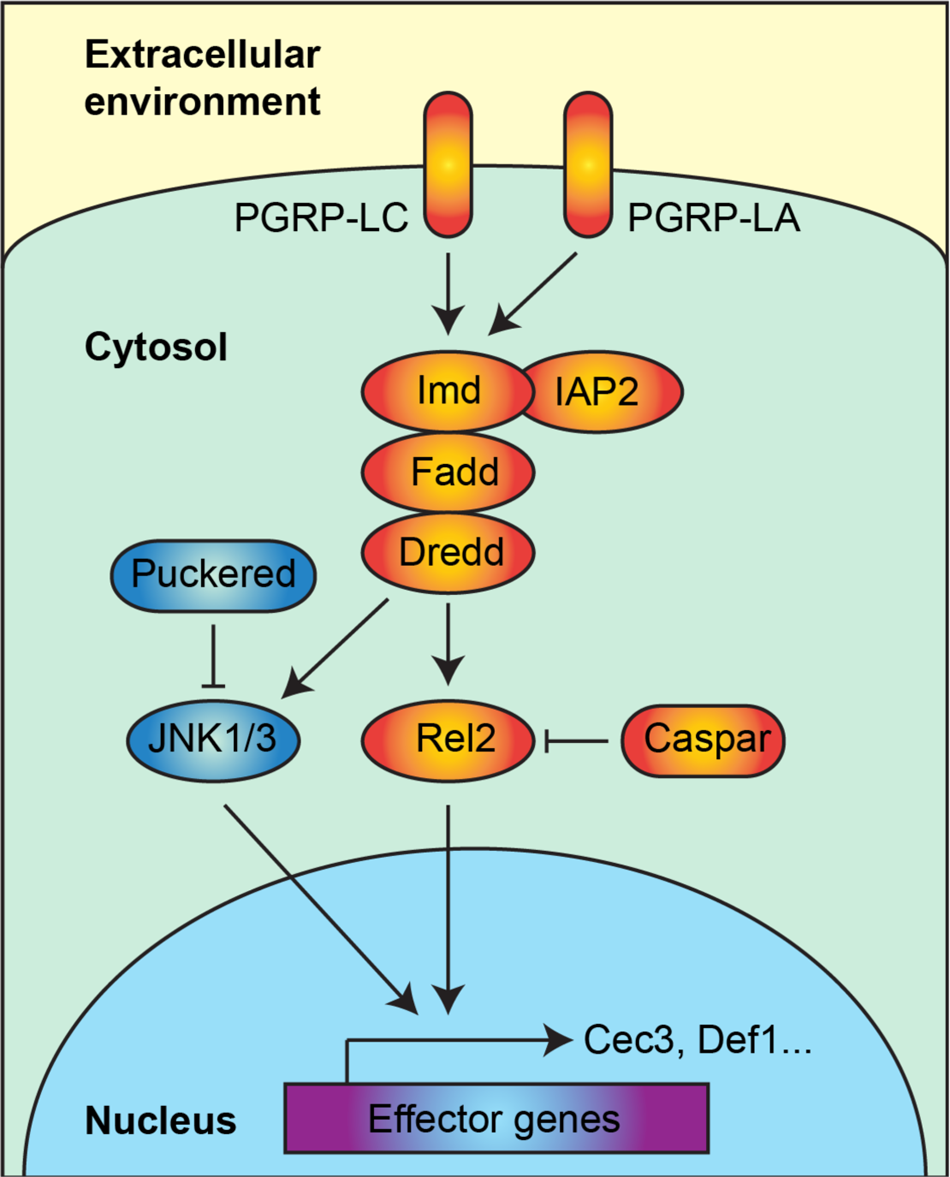
The IMD and interlinked JNK pathways are two of the major immune pathways in mosquitoes. The activation of PGRP-LC triggers the IMD signaling pathway, which includes IAP2, imd, fadd and dredd. This activation dissociates the transcription factor, rel2, from the inhibitor protein, caspar. Rel2 then translocates into the nucleus and induces the production of effectors such as cec3 and def1. Although PGRP-LC is the canonical activator of the IMD pathway, PGRP-LA has also been implicated as an upstream activator of this immune pathway. The IMD pathway bifurcates into the JNK pathway, which activates JNK1 and JNK3, and induces the expression of effector genes. The JNK pathway is negatively regulated by puckered.

**Figure S2.**
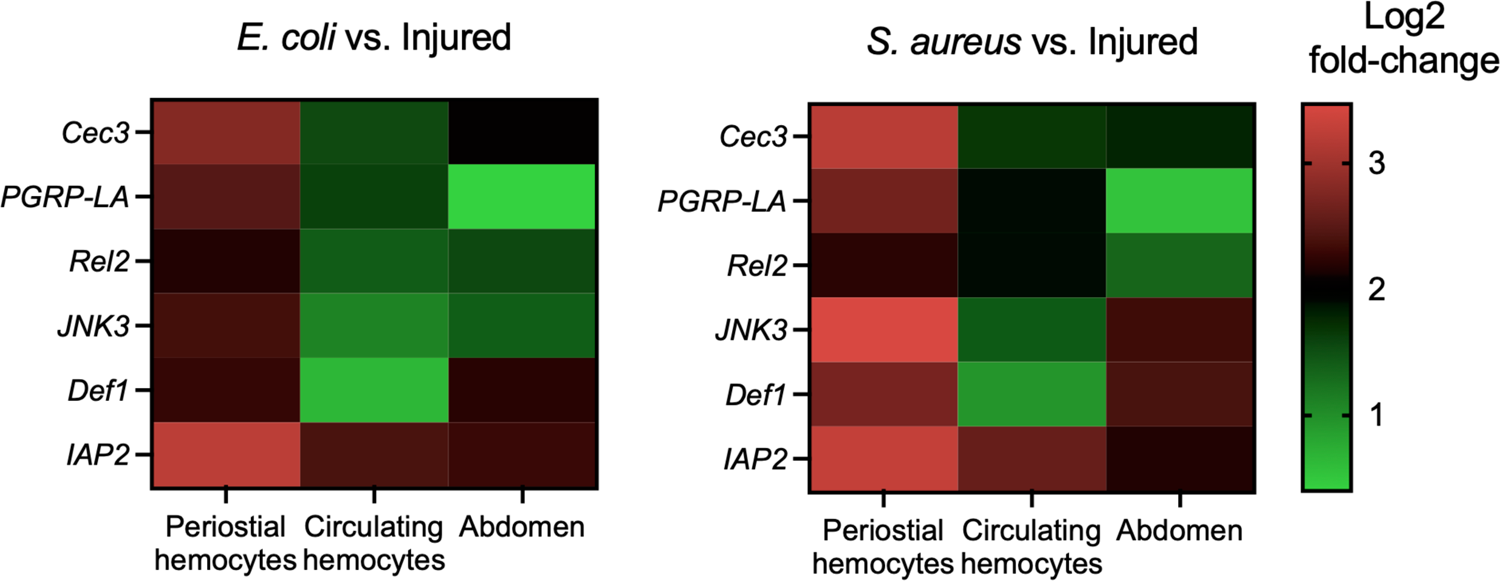
Infection upregulates IMD and JNK pathway genes more highly in the heart and periostial hemocytes than in the circulating hemocytes or the abdomen. Heat maps show that all six IMD and JNK pathway genes identified in our unbiased screen are more highly upregulated after *E. coli* and *S. aureus* infection in the heart and periostial hemocytes than in the other tissues.

**Figure S3.**
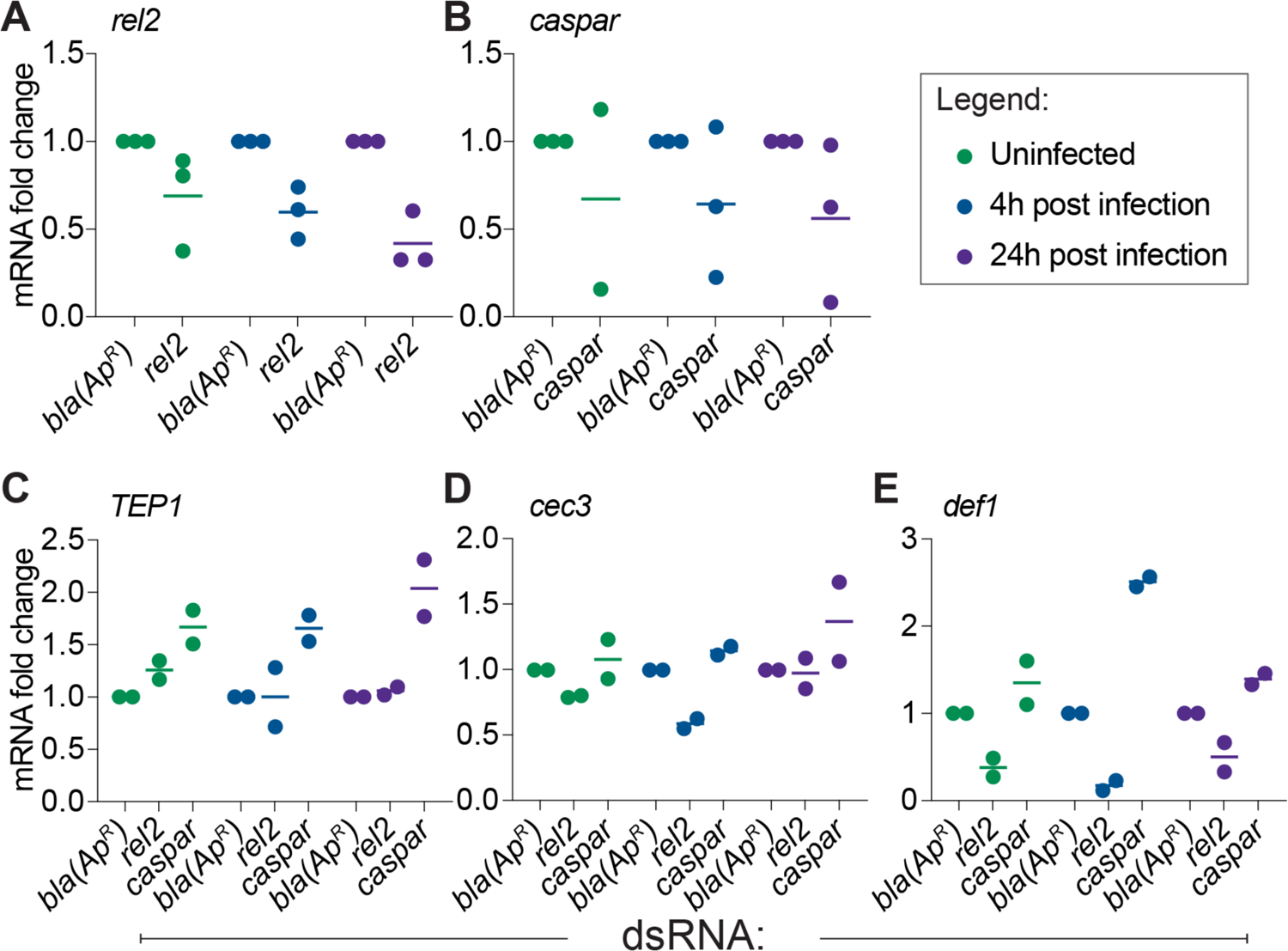
RNA interference-mediated gene silencing of *rel2* and *caspar* reduces their mRNA abundance while altering the mRNA abundance of *TEP1*, *cec3* and *def1*. Relative mRNA abundance of *rel2* (A), *caspar* (B), *TEP1* (C), *cec3* (D), and *def1* (E) in mosquitoes injected ds*bla(Ap^R^)*, ds*rel2* or ds*caspar*. mRNA abundance was measured in mosquitoes that were not infected or had been infected with GFP-*E. coli* for 4 or 24 hr. Horizontal line, mean; Circles, individual biological trials. Data are relative to mRNA abundance in the ds*bla(Ap^R^)* group of that specific trial and treatment.

**Figure S4.**
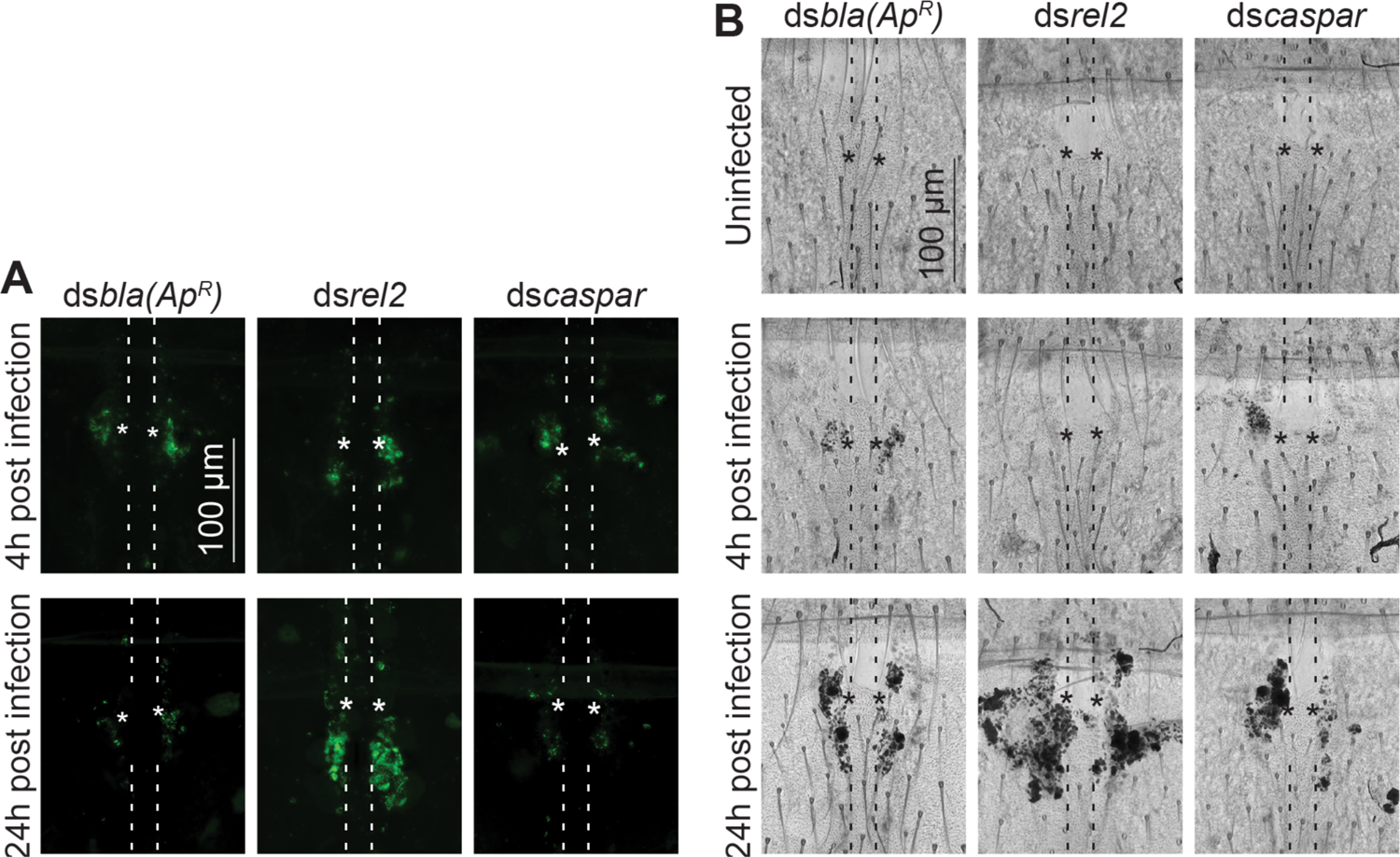
The IMD pathway affects GFP-*E. coli* and melanin accumulation at the periostial regions. (*A*) Fluorescence images show the accumulation of GFP-*E. coli* (green) around a single pair of ostia (asterisks) on a segment of the heart (outlined by dotted lines) of ds*bla(Ap^R^)*, ds*rel2* and ds*caspar* mosquitoes that had been infected with GFP-*E. coli* for 4 or 24 hr. (*B*) Bright field images show the accumulation of melanin (dark deposits) around a single pair of ostia (asterisks) on a segment of the heart (outlined by dotted lines) of ds*bla(Ap^R^)*, ds*rel2* and ds*caspar* mosquitoes that were not infected or had been infected with GFP-*E. coli* for 4 or 24 hr. Anterior is on top. Quantification of all mosquitoes assayed is presented in figures 3A and 3B.

**Figure S5.**
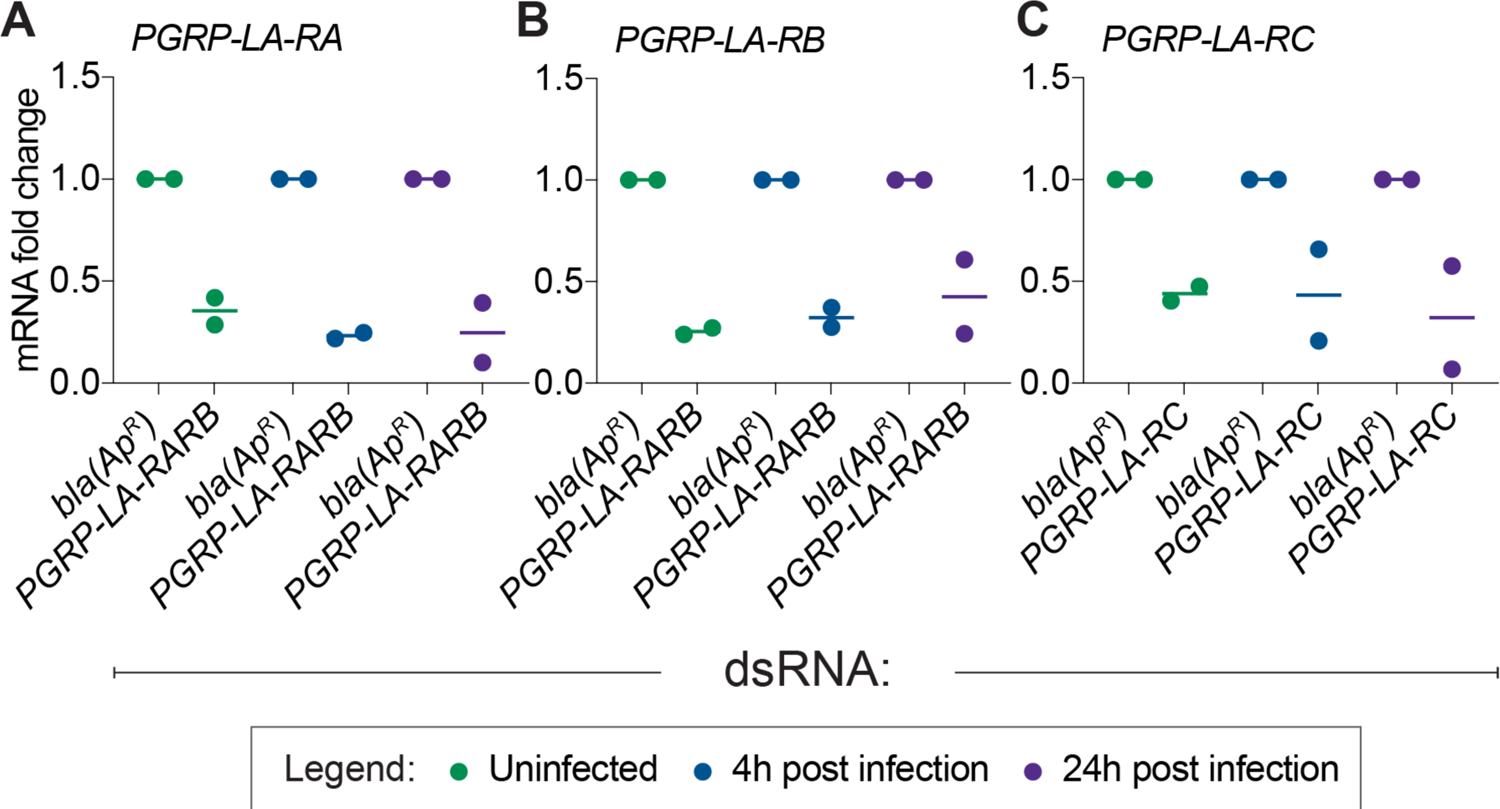
RNA interference-mediated gene silencing of *PGRP-LA* reduces the mRNA abundance of all splice variants. Relative mRNA abundance of *PGRP-LA-RA* (A), *PGRP-LA-RB* (B) and *PGRP-LA-RC* (C) in mosquitoes injected ds*bla(Ap^R^)*, ds*PGRP-LA-RARB* or ds*PGRP-LA-RC*. mRNA abundance was measured in mosquitoes that were not infected or had been infected with GFP-*E. coli* for 4 or 24 hr. Horizontal line, mean; Circles, individual biological trials. Data are relative to mRNA abundance in the ds*bla(Ap^R^)* group of that specific trial and treatment.

**Figure S6.**
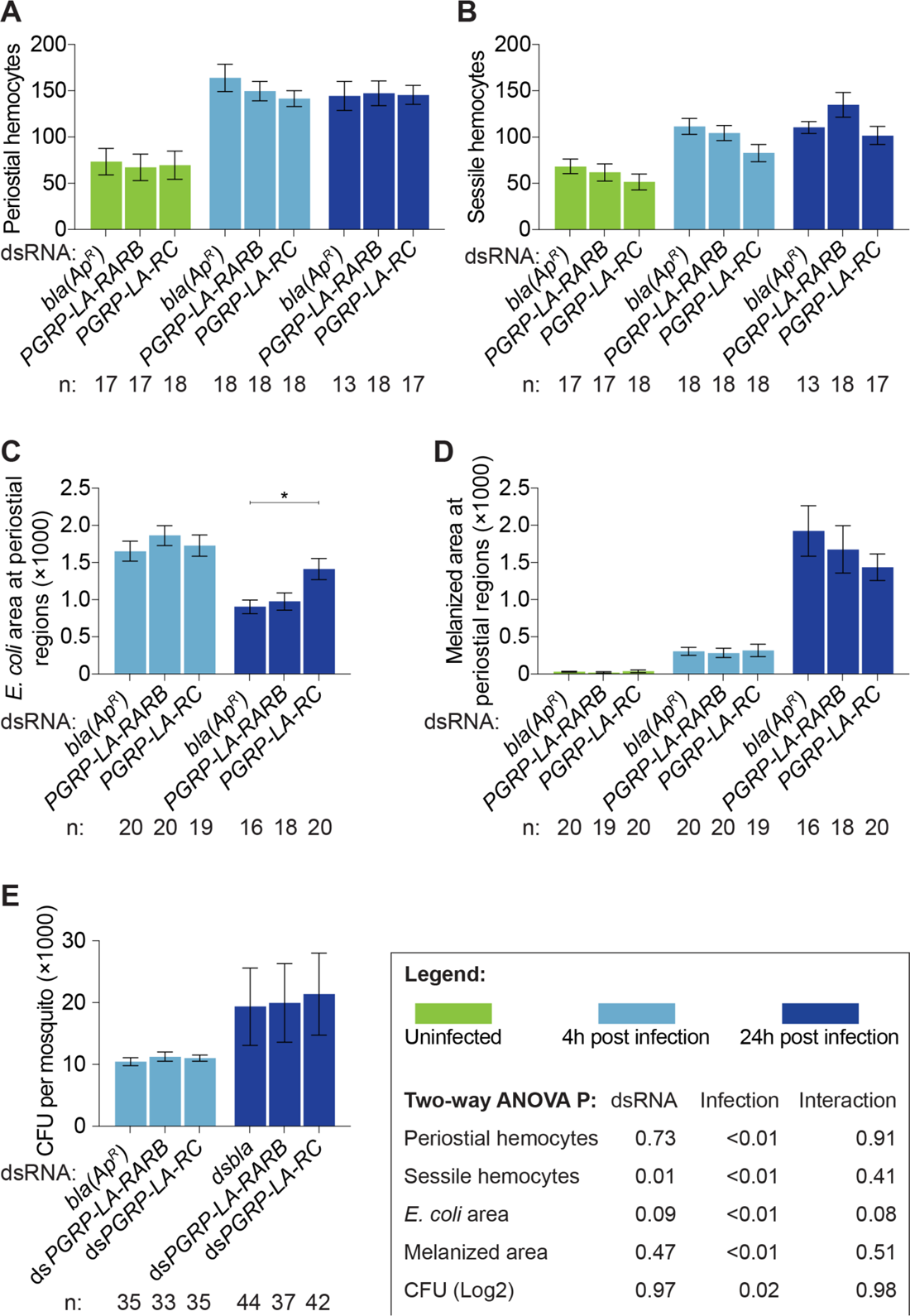
The RC splice form of PGRP-LA regulates phagocytosis by periostial hemocytes. (*A-E*) Graphs for ds*bla(Ap^R^)*, ds*PGRP-LA-RARB* and ds*PGRP-LA-RC* mosquitoes that were not infected or had been infected with GFP-*E. coli* for 4 or 24 hr. The graphs show: (*A*) average number of periostial hemocytes; (*B*) average number of sessile hemocytes outside of the periostial regions in the tergum of abdominal segments 4 and 5; (*C*) pixel area of GFP-*E. coli* in the periostial regions; (*D*) pixel area of melanin in the periostial regions; and (*E*) the systemic GFP-*E. coli* infection intensity. Graphs show the mean and S.E.M. The data were analyzed by two-way ANOVA (bottom box), followed by Dunnett’s multiple comparison test. N indicates sample size. Asterisk in graph indicate post-hoc p < 0.05.

**Figure S7.**
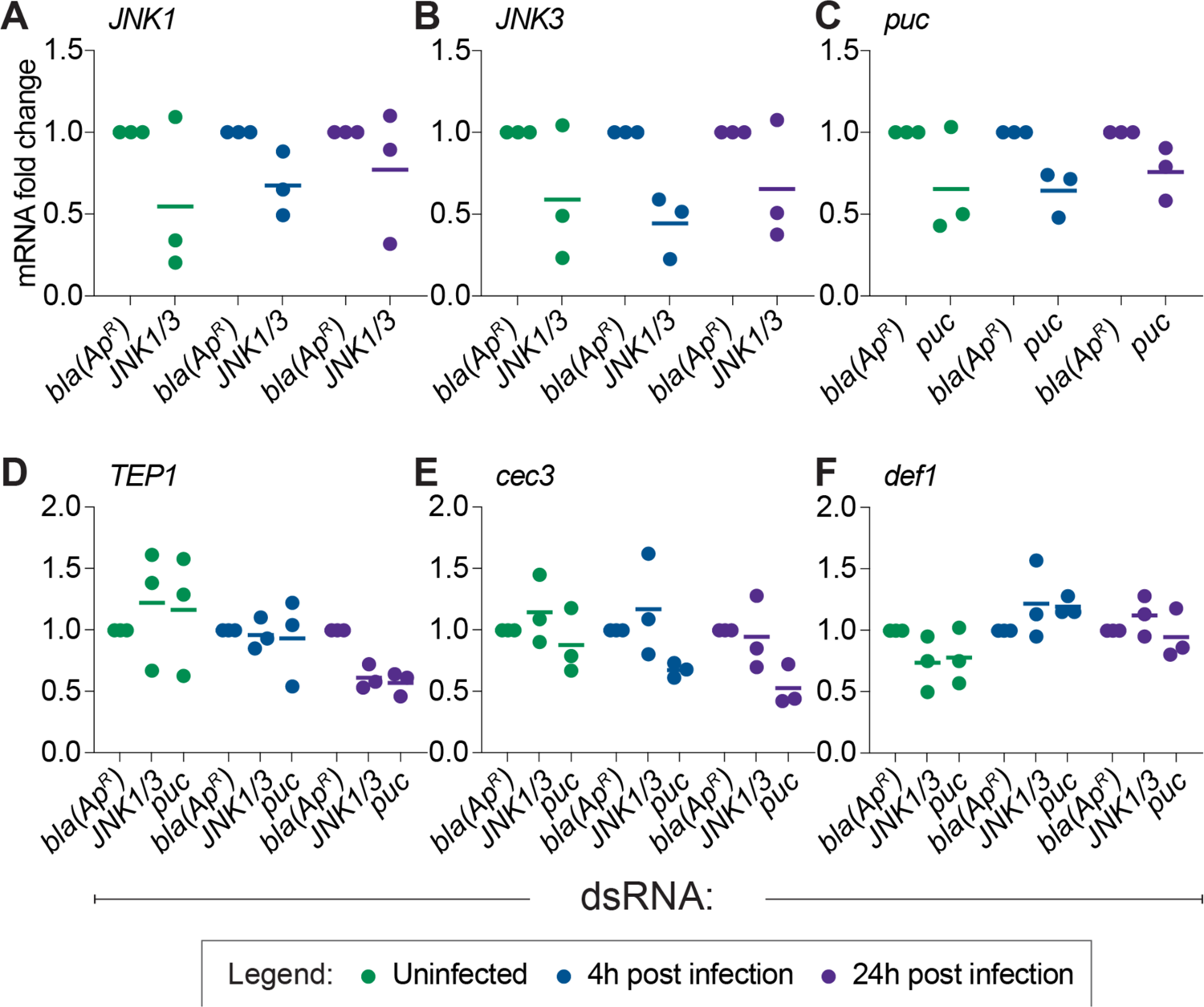
RNA interference-mediated gene silencing of *JNK1*, *JNK3* and *puc* reduces their mRNA abundance while having a limited impact on the mRNA abundance of *TEP1*, *cec3* and *def1*. Relative mRNA abundance of *JNK1* (A), *JNK3* (B), *puc* (C), *TEP1* (D), *cec3* (E), and *def1* (F) in mosquitoes injected ds*bla(Ap^R^),* ds*JNK1/3* or ds*puc*. mRNA abundance was measured in mosquitoes that were not infected or had been infected with GFP-*E. coli* for 4 or 24 hr. Horizontal line, mean; Circles, individual biological trials. Data are relative to mRNA abundance in the ds*bla(Ap^R^)* group of that specific trial and treatment.

**Figure S8.**
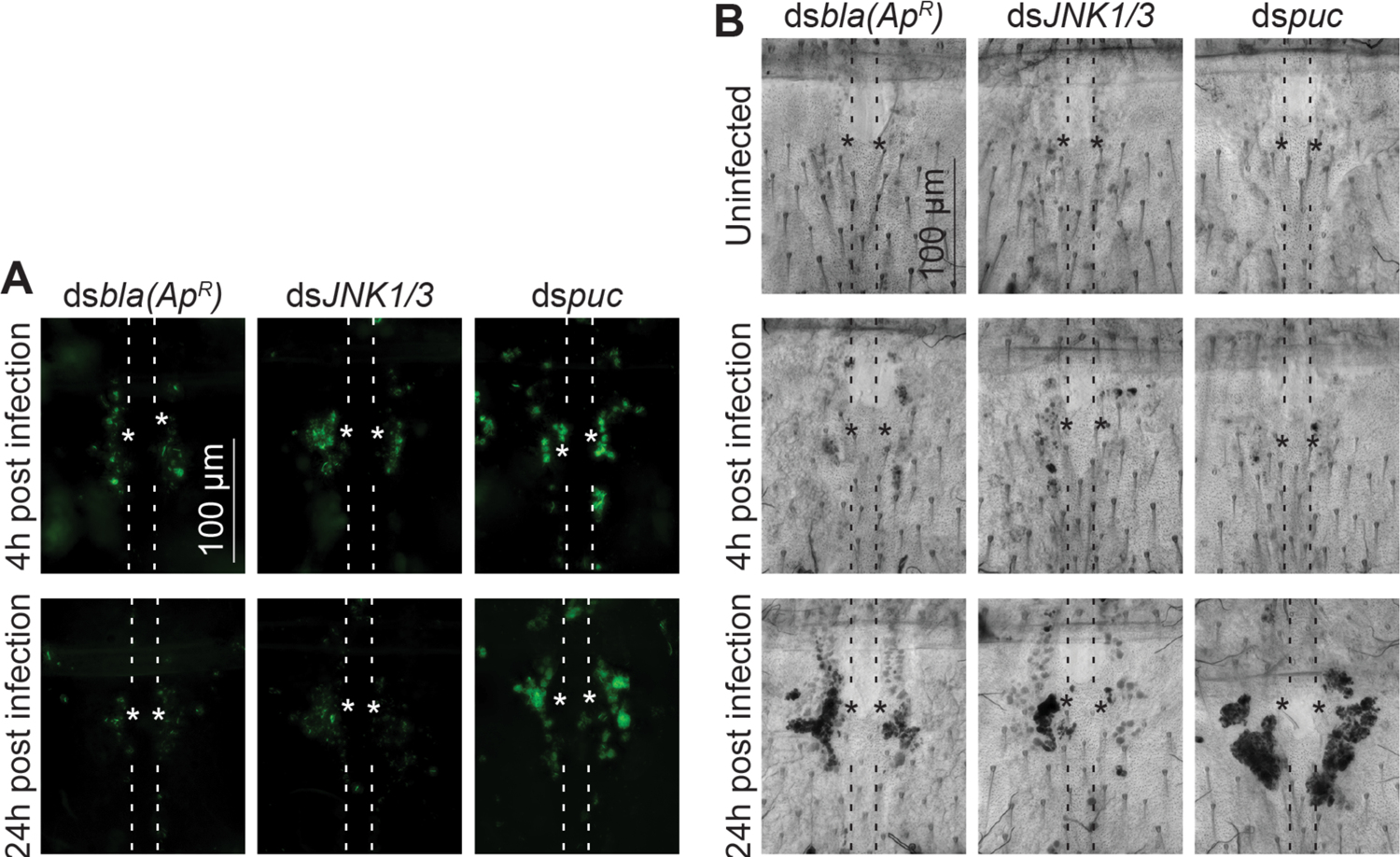
The JNK pathway affects GFP-*E. coli* and melanin accumulation at periostial regions. (*A*) Fluorescence images show the accumulation of GFP-*E. coli* (green) around a single pair of ostia (asterisks) on a segment of the heart (outlined by dotted lines) of ds*bla(Ap^R^)*, ds*JNK1/3* and ds*puc* mosquitoes that had been infected with GFP-*E. coli* for 4 or 24 hr. (*B*) Bright field images show the accumulation of melanin (dark deposits) around a single pair of ostia (asterisks) on a segment of the heart (outlined by dotted lines) of ds*bla(Ap^R^)*, ds*JNK1/3* and ds*puc* mosquitoes that were not infected or had been infected with GFP-*E. coli* for 4 or 24 hr. Anterior is on top. Quantification of all mosquitoes assayed is presented in figures 5A and 5B.

